# Discovery of Phenyl-β-D-glucuronide Medical Function for in Vivo Producing Handheld Gas Sensor Detectable Phenol-like Breath Marker: The Future of Induced Volatolomics in Cancer Risk Pre-warning

**DOI:** 10.1101/2024.02.09.579735

**Authors:** Cuili Xue, Yufei Yan, Huiyan Ni, Chenghan Yi, Yuli Xu, Siyi Xiang, Yuejun Wu, Han Jin, Daxiang Cui

## Abstract

Induced volatolomics provides a promising approach for cancer risk pre-warning. Nevertheless, continues to be questioned bio-compatibility of the synthetic exogenous agents and sophisticated instrument solely detectable isotopes labeled volatile compounds restrained induced volatolomics in clinic application. Here, we report phenyl-β-D-glucuronide as a potential exogenous agent of induced volatolomics. As a nature product, phenyl-β-D-glucuronide demonstrates satisfactory bio-compatibility in pilot study and metabolizes into volatile phenol under the action of tumor micro-environment highly accumulated β-glucuronidase. For conveniently recording breath signal, handheld breath-analyzer based on electrochemical phenol sensor is developed. After administrating healthy or tumor bearing mice with phenyl-β-D-glucuronide, breath signal given by breath-analyzer is verified to be capable of not only predicting various tumor risk with specificity/sensitivity/accuracy above 94.3% in 10 min, but also speculating tumor stage.

## INTRODUCTION

Detecting cancer at early-stage is of utmost important as it significantly increases the chances of successful treatment and survival (*1-3*). Since early-stage tumors are often smaller in size and haven’t spread to other parts of body, surgical removal of the tumor may be sufficient (*4-7*). Basically, early diagnosis improves the chances of complete eradication of cancer or achieving long-term remission, thereby improving the quality and length of life (*2, 8-12*). Nevertheless, cancer in its early stages may not present with any noticeable symptoms (*13-15*), whereas it is normally in the advanced stages if symptoms occur (*15-18*). The absence of warning signs makes it difficult for individuals and even healthcare professionals to identify the presence of cancer. Although, some specific screening tests that available for certain types of cancer have been announced (*19-25*), such as mammograms for breast cancer and colonoscopies for colorectal cancer (*2, 26, 27*), lack of reliable and widely accessible early screening approaches remains concerned issue (*28, 29*).

Recently, directly analysis of specific volatile organic compounds (VOCs) in breath samples has been proved to be a potential and attractive way for cancer diagnosis (*30-35*). Particularly, induced volatolomics essentially improves pre-warning accuracy and thus, further demonstrates its reliability (*36-38*). Principle of the induced volatolomics can be briefly summarized as: in vivo administrating exogenous agents so that to induce some volatile compounds through disease-specific molecular processes; By investigating the level of induced volatile compounds in exhaled breath sample, potential health risk would be easily predicted (*36-39*). However, it should be particularly noted that these so far reported induced volatile compounds are primary in the form of isotope labeled chemicals which can be solely detected via gas chromatography-mass spectrometry (GC-MS) (*39*). Hence, it is hardly for customers to predict their cancer risk by simply implementing breath analysis at home. Besides, bio-compatibility (e.g. toxicology) of previous reported exogenous agents still remains questionable (*39, 40*). In sum, these aforementioned challenge factors restrict clinic application of the induced volatolomics in early cancer detection.

Potential approach to overcome these challenges is to develop an exogenous agent that meets the following criteria: i) to avoid any unclear adverse effects, natural products or chemicals extracted from natural source are preferred for the exogenous agent; ii) non-toxic or negligible toxicity is required for the trace level organic volatile compounds that produced during exogenous agent in vivo metabolism. Meanwhile, there should be distinct difference in the concentration of induced volatile compound in exhaled breath between healthy individuals and cancer patients; iii) to enable users evaluating their cancer risk at home, the induced volatile compound should be easily detectable with cheap gas sensors. Herein, we report on the discovery of adopting phenyl-β-D-glucuronide, a natural product found in Glycine max metabolite (*41*), to be a satisfactory exogenous agent for induced volatolomics-based cancer detection (Figure 1a). Under the enzymatic action of β-glucuronidase, phenyl-β-D-glucuronide is expected to be metabolized into trace amount of phenol which may finally appear in exhaled breath (Figure 1 b). Note that β-glucuronidase accumulates selectively in the micro-environment of numerous solid tumors while this enzyme is essentially located inside the cells in healthy tissues (*36*), therefore, after taken phenyl-β-D-glucuronide for a while (within 2 h), significant difference in the level of phenol is speculated to be found in the breath samples that derived from health individuals or cancer group. Moreover, chemical active phenol allows us to develop handheld breath-analyzer that based on an electrochemical gas sensor to track its concentration variation which disclosing bright future of implementing the self-diagnosis at home (Figure 1 c). Herein, we systematically investigate the practicability of adopting phenyl-β-D-glucuronide to early cancer screening. In particular, a portable sensing device that integrated a highly selective electrochemical phenol sensor is developed to realize the convenient non-invasive pre-warning.

**Fig. 1.**
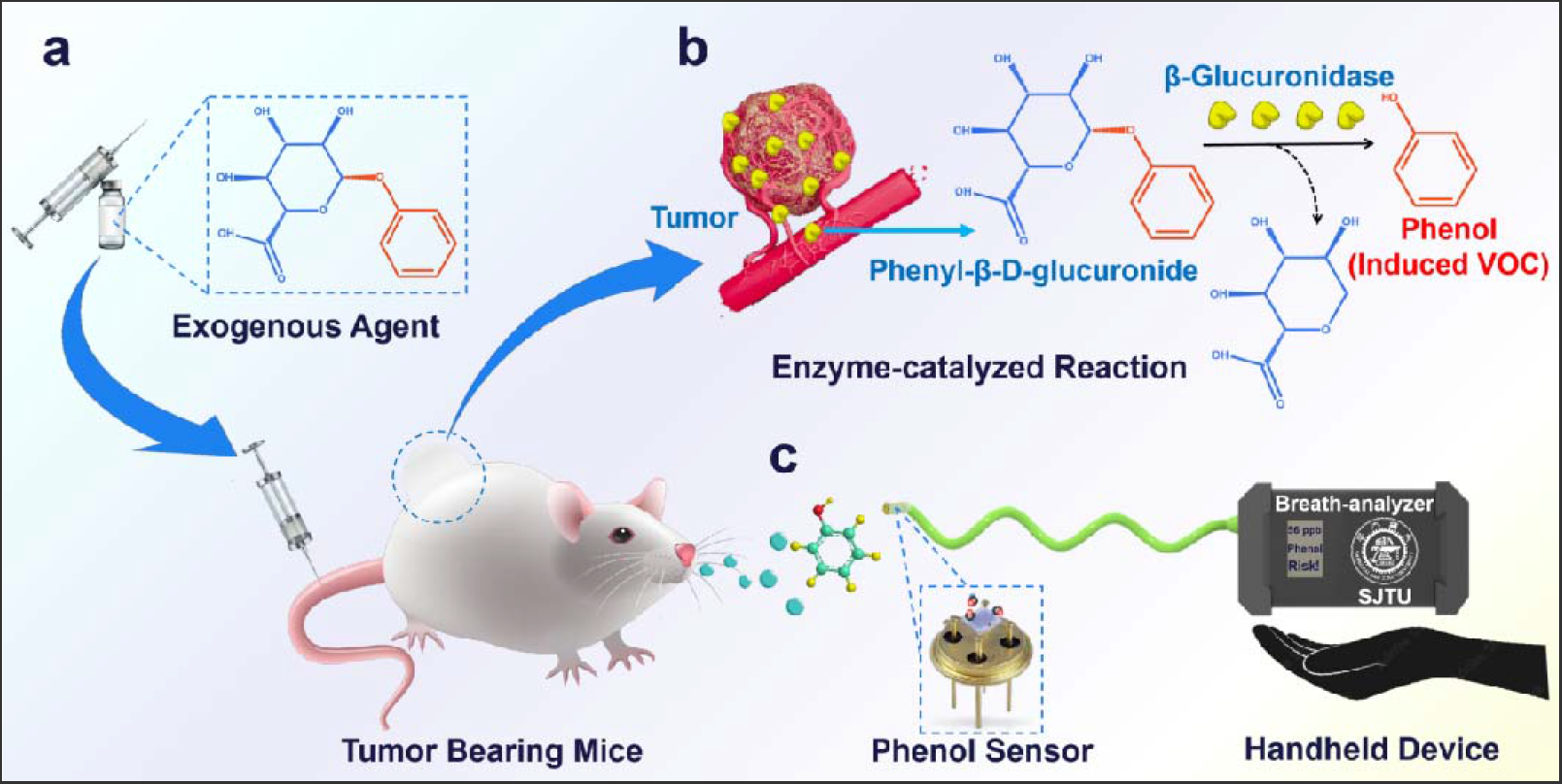
Illustration of the strategy: **a**, Phenyl-β-D-glucuronide is employed as the exogenous agent and delivered to tumor location by means of intravenous injection and blood circulation; **b**, After closing to tumor, phenyl-β-D-glucuronide starts to release phenol (induced VOC) under the enzyme action of β-glucuronidase; **c**, Induced phenol is expected to be appeared in exhaled breath and its level will be further monitored via a handheld breath-analyzer. The risk of suffering cancer can be easily predicted based on the level of phenol in breath samples.

## RESULTS AND DISCUSSION

### Exogenous agent synthesis and enzymatic products characterization

Although phenyl-β-D-glucuronide can be directly purified from the metabolite of Glycine max, we still report its synthetic method for future mass production (Figure 2a). The details are presented in Supplementary Material and Methods section. In brevity, the synthesis of phenyl-β-D-glucuronide commenced with methyl 1,2,3,4-tetra-O-acetyl-β-D-glucuronate (raw material 1), the raw material 1 treated with phenol using a tin(iv) chloride catalyst at room temperature in dichloromethane (DCM) for 16 hours and provide intermediate 2 (verified by infrared spectra (IR), nuclear magnetic resonance spectroscopy spectra (NMR) and mass spectra (MS) that shown in Supplementary Figures 1 and 2) in 41.8% yield. Next, the Intermediate 2 is hydrolyzation with sodium Methoxide at room temperature in MeOH for 2 hours and provide final phenyl-β-D-glucuronide (product 3) in 35.1% yield. Chemical structure of the synthesized phenyl-β-D-glucuronide is further verified by IR (Figure 2b, (neat, cm^−1^): 3502, 3456, 2900, 1743, 1600, 1492, 1415, 1226, 1080, 752, 690, 636) and NMR (Figure 2c, ^1^H NMR (400 MHz, D_2_O) δ 7.44 (t, *J* = 7.7 Hz, 2H), 7.24 – 7.15 (m, 3H), 5.22 – 5.16 (m, 1H), 4.03 (dd, *J* = 6.4, 3.7 Hz, 1H), 3.72 – 3.61 (m, 3H) and Supplementary Figure 3a) as well as MS spectra (Supplementary Figure 3b). All these characteristic peaks are consistent with that of phenyl-β-D-glucuronide (CAS: 17685-05-1), suggesting the success of obtaining high-purity phenyl-β-D-glucuronide.

**Fig. 2.**
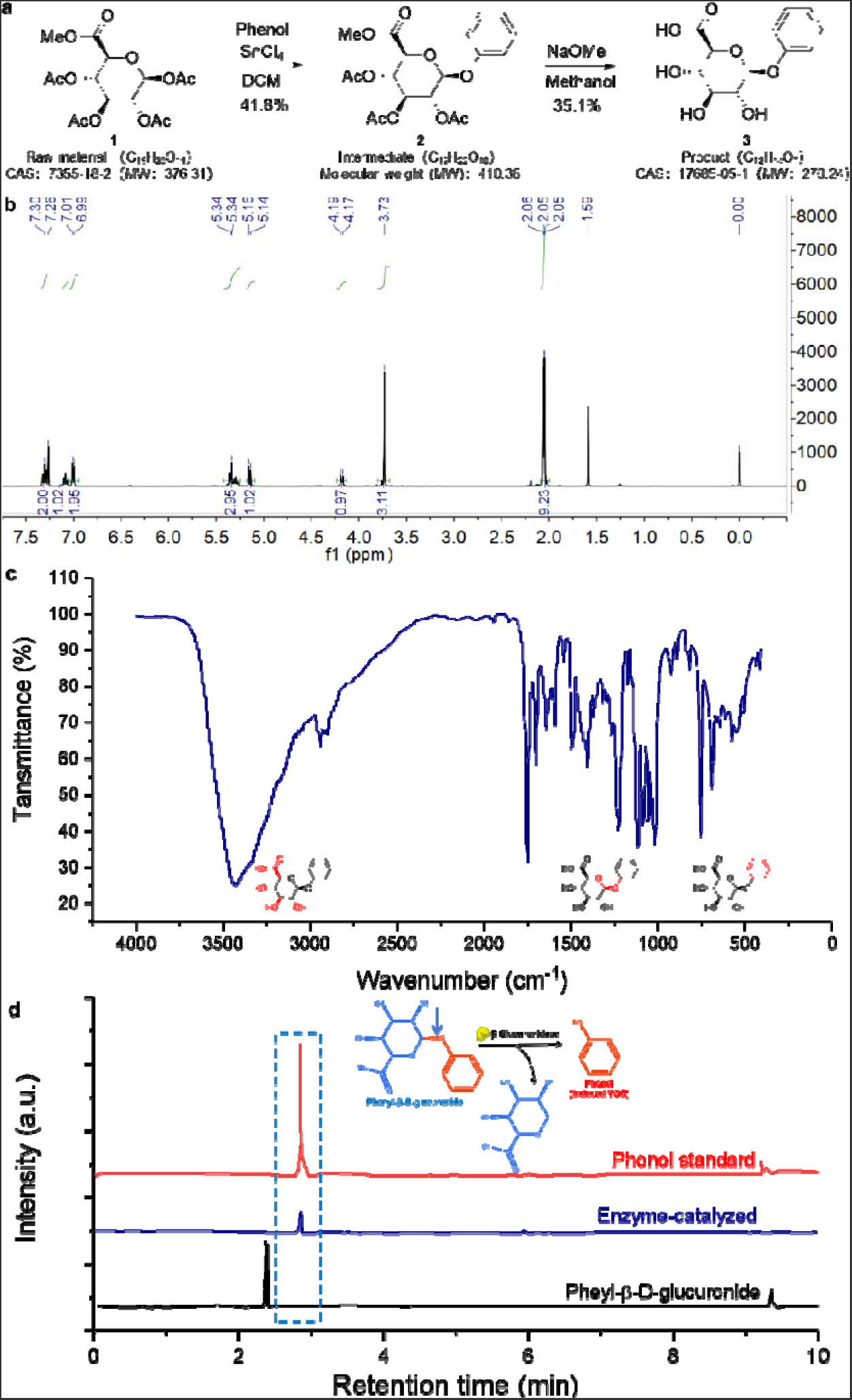
Phenyl-β-D-glucuronide synthesis steps and verification of the induced phenol during enzymatic reaction. **a**, Steps of synthesizing phenyl-β-D-glucuronide; **b**, HNMR (Hydrogen NMR) and **c**, IR spectra of the phenyl-β-D-glucuronide product that obtained via the proposed synthesis approach; **d**, LC spectra of the synthesized phenyl-β-D-glucuronide and its enzymatic hydrolysis product that catalyzed by β-glucuronidase, as well as phenol standard.

Figure 2d demonstrates the liquid chromatography (LC) spectra for the samples of synthesized phenyl-β-D-glucuronide and its enzymatic hydrolysis products that catalyzed by β-glucuronidase. Meanwhile, LC spectra of phenol standard is also provided for comparison. According to the LC spectra, it is clear that β-glucuronidase effectively promotes the hydrolysis reaction and yielded abundant of phenol. To confirm whether the induced phenol can appear in gas phase or not at room temperature, volatile compounds in upper gas phase of all the aforementioned samples are also collected and analyzed through gas chromatography (GC). Supplementary Figure 4 implies that volatilized phenol is solely found in upper gas phase of the sample containing enzymatic hydrolysis products. These important results speculate the potential of in-vivo generating induced phenol for the tumor bearing objects that have taken phenyl-β-D-glucuronide like exogenous agent, particularly, the induced phenol is highly possible to be found in exhaled breath.

### Toxicology and pharmacokinetic parameters of phenyl-β-D-glucuronide like exogenous agent

It is generally known that bio-compatibility of the exogenous agent is the primary consideration for future clinical application. In this case, it is highly valuable to confirm whether phenyl-β-D-glucuronide, reported as a natural product, has good bio-compatibility. Consequently, toxicity of the chemical in vitro is thoroughly studied and presented in Figure 3. Initially, cell viability of phenyl-β-D-glucuronide solution (diluted with phosphate buffer saline, PBS solution) is verified on human gastric mucosal epithelial cells by standard cell-counting kit 8 (CCK-8) analysis. As shown in Figure 3a, the cell viability remained at 92.3 % even the exogenous agent concentration up to 500 μg/mL, indicating negligible cytotoxicity against cells incubated with phenyl-β-D-glucuronide soluton. Then, body weight of the healthy mice that dosed without (control group) or with 200 μg/kg phenyl-β-D-glucuronide were compared. The similarity of body weight for all groups at different measuring period implies that dosing the phenyl-β-D-glucuronide did not cause any notable toxic side-effects (Figure 3b). Hematoxylin Eosin (H&E) stained sections of major organs (heart, liver, spleen, lung, kidney) of phenyl-β-D-glucuronide treated healthy and tumor bearing mice also exhibit no obvious pathological changes (Figure 3c). In addition, animal testing reports of acute systemic toxicity test and the maximization test for sensitization reveal that extract solution of phenyl-β-D-glucuronide did not cause acute systemic toxicity and/or skin allergic reaction (Supplementary reports for acute systemic toxicity test and the maximization dosage test). In conclusion, all these aforementioned preliminary toxicology test results validate good bio-compatibility of the phenyl-β-D-glucuronide like exogenous agent.

**Fig. 3.**
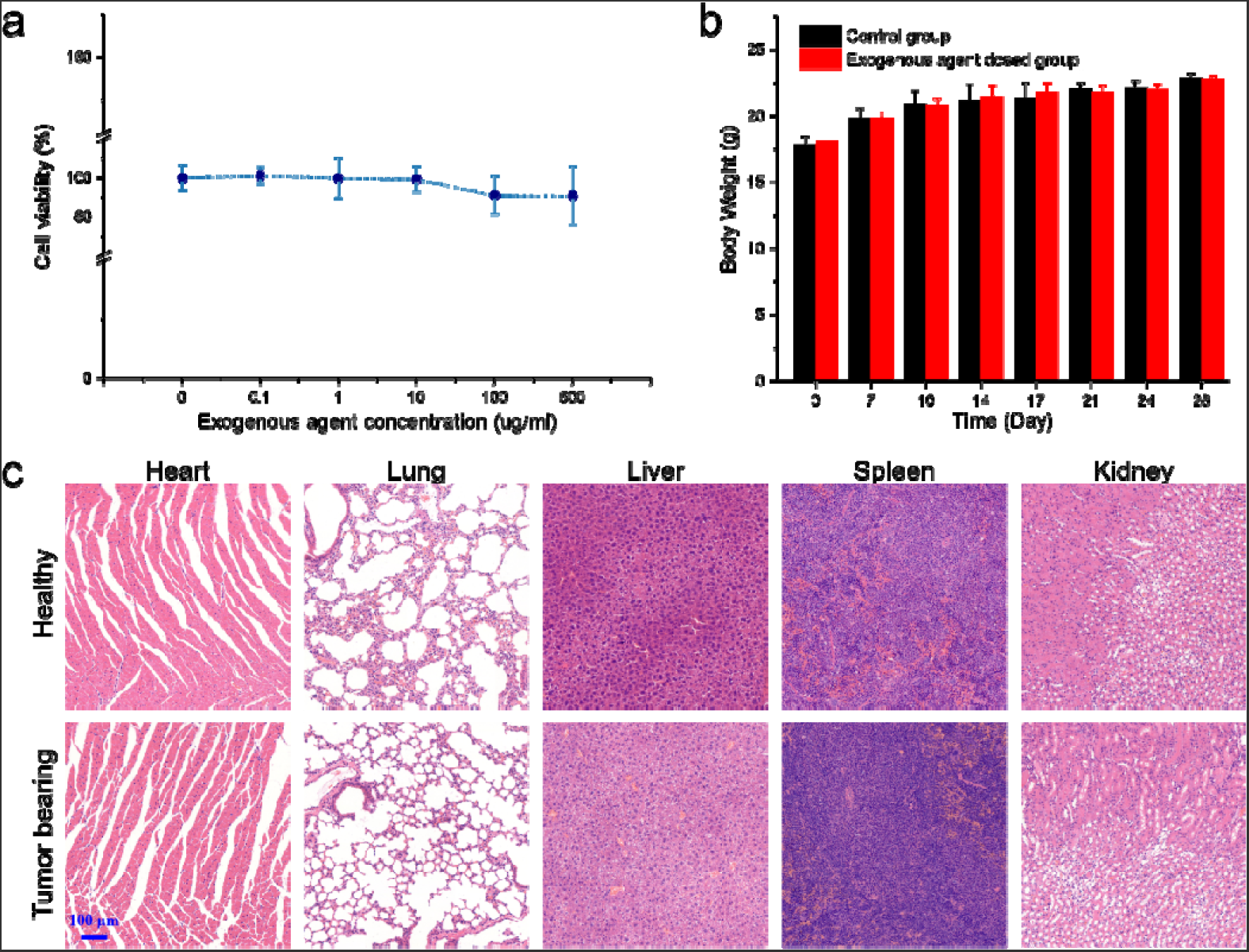
Bio-compatibility of phenyl-β-D-glucuronide and its pharmacokinetic parameters. **a**, Cell viability of human gastric mucosal epithelial cell lines incubated with different concentration phenyl-β-D-glucuronide. **b**, Comparison of body weight for the healthy mice administrated without (control group) or with 200 μg/kg phenyl-β-D-glucuronide at different cultivation days. **c**, H&E stained the heart, lung, liver, spleen and kidney specimens from the healthy and tumor bearing mice that have been treated with phenyl-β-D-glucuronide for 4 times.

### In vivo investigation of phenyl-β-D-glucuronide induced breath signal

Medical function of phenyl-β-D-glucuronide in cancer risk pre-warning is thoroughly assessed by investigating the concentration difference of the induced phenol in breath sample that exhaled from tumor bearing mice or healthy counterparts. Before starting the investigation, technique parameters such as phenyl-β-D-glucuronide dosage, temperature of the mice incubator during sampling and the sampling time after administrating exogenous agent are optimized. Supplementary Figure 5 indicates that the optimal exogenous agent dosage, incubator temperature and sampling time are 200 μg/kg, 28 °C and 1 h, respectively. Based on these technique parameters, breath sample is collected and analyzed. In briefly, after tail vein injecting phenyl-β-D-glucuronide for 1 h, breath samples exhaled from health or tumor bearing group (e.g. gastric tumor bearing mice) are collected and further analyzed with gas chromatography-mass spectrometry (GC-MS), as illustrated in Figure 4a. At the initial stage, phenol level that derived from different breath sample is simply estimated by directly comparing the GC-MS peak intensity. According to the statistical data shown in Figure 4b, it can be concluded that prior to the tumor volume essentially increase, minor difference for the induced phenol level is found for in the breath samples derived from healthy or tumor bearing group. Nevertheless, apparent increment in the induced phenol level can be observed after tumor significantly growth. In contrast, the induced phenol exhaled from healthy mice almost maintains as a constant level, regardless of the cultivation time. Impressively, level of the induced phenol increases with the increasing of tumor volume and linear relationship is proved between these two parameters (Figure 4c, d).

**Fig. 4.**
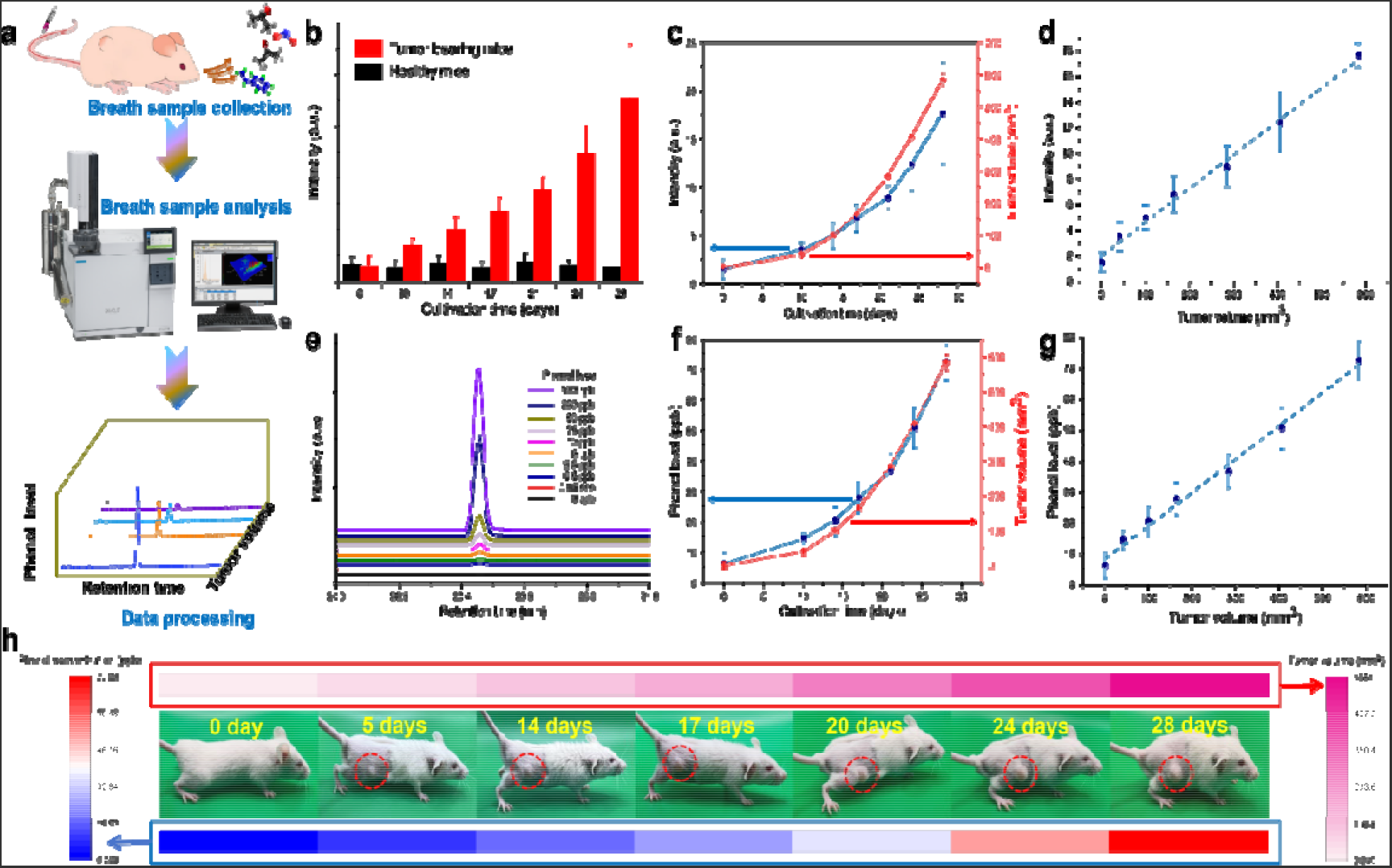
Feasibility of using phenyl-β-D-glucuronide to trigger a rapid breath readout of potential gastric cancer risk. **a**, Flowchart of investigating phenyl-β-D-glucuronide induced breath signal. **b**, Amount of induced phenol in breath sample that released by healthy mice (black bars; n=23) and mice with gastric cancer subcutaneous xenograft (tumor bearing mice, red bar, n=23) at different cultivation days. **c, d**, Potential relationship between the exhaled phenol level and tumor volume. **e**, Phenol level calibration. **f, g**. Function relationship of induced phenol level in exhaled sample on tumor volume. **h**, Phonographic image of the tumor bearing mice cultivated within 28 days and the induced phenol level in exhaled sample vs. tumor volume.

To implement quantitative analysis, the induced phenol level in diverse breath sample is calibrated (Figure 4e and Supplementary Figure 6). After processing the data with standard curve method (Figure 4f-h), it is reasonable to deduce that the induced phenol level exhaled from healthy mice is around 7.2 ppb while that of the level in the breath sample of tumor bearing mice ranges from 6.8 ppb (tumor volume ≈ 0 mm^3^) to 72.6 ppb (tumor volume ≈ 583.8 mm^3^). For instance, the induced phenol level exhaled form healthy mice and tumor bearing mice that cultivated for 10 days (tumor volume ≈ 40.6 mm^3^) is roughly estimated to be 6.3 ppb and 14.6 ppb. Meanwhile, functional relationship of the induced phenol level and tumor volume is described in equation (1). Conclusively, administration of phenyl-β-D-glucuronide could trigger a rapid breath readout of potential gastric cancer risk via analyzing the induced phenol level.

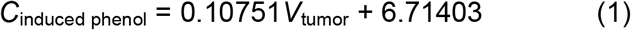

Where *C*_induced phenol_ (ppb) and *V*_tumor_ (mm^3^) represent the induced phenol level and volume of subcutaneous tumor, respectively.

### Cancer risk prediction via a handheld breath-analyzer

In addition to verifying the feasibility of utilizing phenyl-β-D-glucuronide for volatolomics-based cancer risk pre-warning, developing a low cost and easy to operate device for breath markers detection is also an urgent issue that needs to be addressed. Herein, handheld breath-analyzer that based on a miniaturized electrochemical gas sensor is particularly proposed. For the purpose of high performance tracking the phenol-based breath marker that derived from phenyl-β-D-glucuronide metabolism, initially, gas adsorption capability of various metallic oxides is compared to screen out the phenol adsorption preferred sensing material (Supplementary Figure 7). Note that volatile compounds (VOCs, e.g. acetone, styrene, hexanal, ethanol, n-butanol, ethylbenzene) that can be primary found in exhaled breath are selected as the interference gases in this study (Figure 5a and Supplementary Figure 7). As shown in Supplementary Figure 7 that among all the examined metallic oxides, relative larger frequency shift for phenol is solely observed for Cr_2_O_3_. Since large frequency shift suggests the high affinity to specific gas species, it is reasonable to deduce that Cr_2_O_3_ prefers to capture phenol molecular in comparison with that of other examined VOCs. This unique physical and chemical properties enable phenol to be highly selective adsorbed on the surface of Cr_2_O_3_. Due to the fact that phenol preferred adsorption is one of the crucial step to realize the selectivit, Cr_2_O_3_ is believed to be a good candidate for sensor fabrication. Then a compact yttria stabilized zirconia (YSZ) based gas sensor that comprised of Cr_2_O_3_ sensing electrode (SE) and Mn-based reference electrode (RE) is designed, and its operating parameters (i.e. fabrication and operating temperature) are also optimized (Supplementary Figure 8). Optimal response magnitude to 0.6 ppm phenol is demonstrated at the fabrication and operating temperature of 1050 °C and 450 °C, respectively. After that, sensing behavior of the sensor is thoroughly characterized. In summary, the developed sensor gives high specificity to phenol against other examined VOCs (Figure 5a) and a linear relationship between the response signal and phenol concentration is confirmed (Figure 5b, c). Besides, 90% response/recovery time of the sensor is roughly estimated to be 37 s/41 s (Supplementary Figure 9a). Such acceptable response/recovery speed would be beneficial to the rapid self-monitoring of user. Furthermore, the sensor shows satisfactory response stability and moisture resistance when detecting phenol (Supplementary Figure 9b, c). The impressive sensing characteristics imply the practicability of the sensor in tracking trace amount of induced phenol in breath sample.

**Fig. 5.**
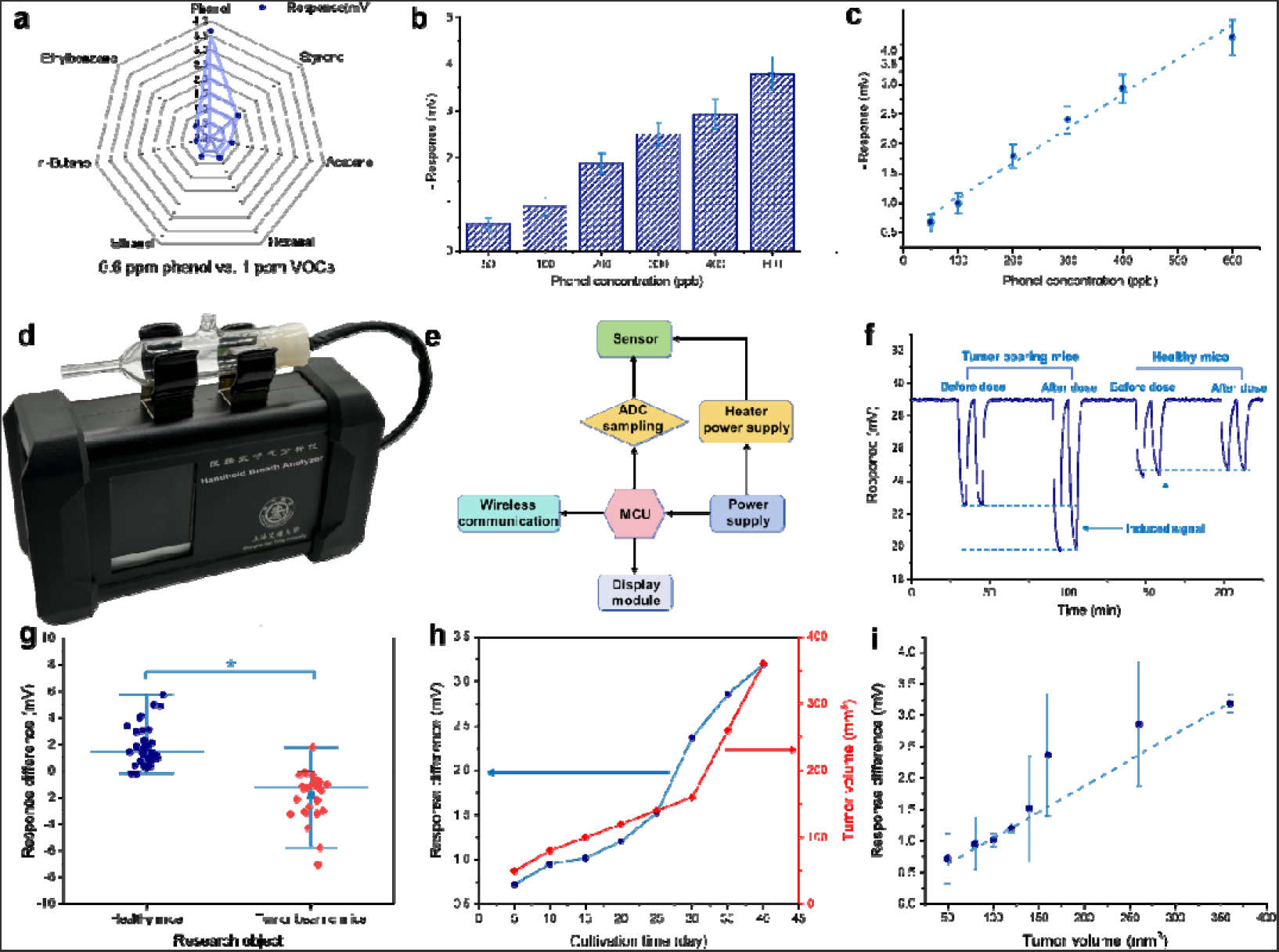
Pre-warning of cancer risk through the proposed breath-analyzer. **a**, Comparison of response signal to various studied gas species. **b**, Step response of the sensor to phenol in the concentration of 50-600 ppb. **c**, Dependence of the response signal for the sensor on the level of phenol, within the examined concentration. **d**, Photograph and **e**, operating diagram of the handheld breath-analyzer; **f**, Variation of the breath signal measured before or at 1h after dosing phenyl-β-D-glucuronide. Apparent shifting in the signal is found for the breath samples derived from tumor bearing mice, confirming the practicability of predicting cancer risk via the breath-analyzer. **g**, Comparison of the response difference for the healthy and tumor bearing mice. Circle and diamond symbols indicate healthy and gastric tumor bearing mice, respectively (mean□±□s.d., N=□2 independent experiments indicated in shades, n=□32 or 33 mice per group, two-tailed Mann–Whitney test, ^***^P=□0.0005). **h**, Variation of the response difference and tumor volume on cultivation time. **i**, Dependence of response difference on the tumor volume. Approximate linear relationship between the response difference and tumor volume indicates the potential of predicting the cancer stage through the breath signal difference given by breath-analyzer.

A handheld breath-analyzer that integrated the aforementioned YSZ-based electrochemical phenol sensor is proposed to implement the expected non-invasive cancer pre-warning (Figure 5d, Supplementary Figure 10) and its overall operating diagram are summarized in Figure 5e. Breath samples for the healthy mice or tumor bearing mice are collected and phenol concentration in all collected breath samples is examined via the breath-analyzer. Prior to investigating the variation of breath signal induced by phenyl-β-D-glucuronide, impact of the solvent (i.e. PBS) that utilized for preparing phenyl-β-D-glucuronide solution at certain concentration is studied and negligible impact on the breath signal is observed, indicating the variation of breath signal will be solely contributed by the administration of phenyl-β-D-glucuronide (Supplementary Figure 11). The breath signal that recorded by the breath-analyzer is analyzed for the mice before or after dosing phenyl-β-D-glucuronide. After dosing phenyl-β-D-glucuronide, apparent increment in breath signal is solely observed for breath samples derived from gastric tumor bearing mice (Figure 5f). The obvious distinction in response difference (response difference = “breath signal measured before dosing phenyl-β-D-glucuronide” - “breath signal measured 1h after dosing phenyl-β-D-glucuronide”) between the breath samples derived from healthy mice and tumor bearing mice demonstrates the ability of phenyl-β-D-glucuronide assisted breath analysis technology in predicting potential cancer risk (Figure 5g, P=□0.0005). Particularly, as the tumor volume increases, response difference will continuously increase (Figure 5h). Meanwhile, an approximate linear relationship between the response difference and the tumor volume is further confirmed (Figure 5i) and the functional relationship is summarized in equation (2), suggesting that it is highly possible to deduce the gastric cancer stage through judging the response signal given by the breath-analyzer.

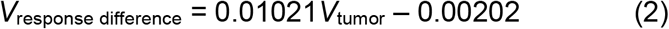

Where *V*_response difference_ (mV) and *V*_tumor_ (mm^3^) represent the response difference given by breath analyzer and volume of subcutaneous tumor, respectively. Additionally, accuracy, specificity and sensitive of the proposed approach are around 95.4%, 94.3% and 96.7%, respectively (confirmed by double blind test with n = 65). Taken together, these pilot results illustrate the success for the utility of phenyl-β-D-glucuronide like exogenous agent and handheld breath-analyzer platform for assessing the cancer risk which could be easily realized at home.

## SUMMARY

Here, we report the potential medical function of phenyl-β-D-glucuronide for non-invasive pre-wanring of tumors. Particularly, we provide synthetic route for future mass production of phenyl-β-D-glucuronidelike exogenous agent and disclose that phenyl-β-D-glucuronide like exogenous agent can be enzymatically converted by β-glucuronidase, an extracellular enzyme secreted by solid tumours, into phenol which is then detectable in exhaled breath. Animal test verified the significantly increase in phenol level for the breath sample derived from tumor bearing mice that subjected to phenyl-β-D-glucuronide administration, and thus offering a rapid, sensitive and non-invasive approach for pre-warning of potential cancer risk. Moreover, as a nature product of Glycine max metabolite, phenyl-β-D-glucuronide shows satisfactory bio-compatibility in pilot study.

We also develop a handheld breath-analyzer that based on a miniaturized YSZ-based gas sensor for high-performance detecting the induced phenol in breath. It is verified that breath signal given by the breath-analyzer is not only helpful to predicting a variety of potential cancer risk, but could also be used to speculate tumor stage. We envision that future clinic application of the phenyl-β-D-glucuronide like tumor probe and handheld breath-analyzer platform may offer the opportunity of self-assessing cancer risk at home.

It should be additionally noted that although we have conducted preliminary studies on the bio-compatibility of phenyl-β-D-glucuronide, its in-vivo toxicity should be particularly thoroughly characterized due to the chemical has never been used in human clinical trials. Besides, tumor bearing mice is solely selected as the tumor model in our pilot study, consequently, similar investigation should be further implemented with in situ tumor model in future for systematically validating the practicability of the proposed technology.

## Supporting information

Supplmental Materaisl and Methods

Supplementary reports for acute systemic toxicity test.

Supplementary reports for the maximization dosage test

## ACKNOWLEDGMENTS

We acknowledge the Analysis Testing Center at Shanghai Jiao Tong University (SJTU) for GC-MS, IR and NMR technical support, as well as the Center for Advanced Electronic Materials and Devices (AEMD) at SJTU for sensor device fabrication technical support. We thank S. S. Cui for providing suggestion of NMR results analysis and Y. L. Liu giving help for tumor bearing mice cultivation. We also gratefully acknowledge the financial support for this research from the National Natural Science Foundation of China (Grant No. 62227815).

## Author contributions

C. L. X. conducted the research of exogenous agent synthesis and enzymatic products characterization as well as in vivo investigation of phenyl-β-D-glucuronide induced breath signal. Y. F. Y. designed and fabricated the handheld breath-analyzer and implemented animal test. H. Y. N. assisted with the exogenous agent synthesis and characterization. C. H. Y. and Y. L. X. fabricated the YSZ-based phenol sensor and evaluated its sensing characteristics. S.Y. X. and Y. J. W. assisted with the circuit board design of the breath-analyzer. H. J. and D. X. C. designed all experiments and supervised the research. H. J. wrote the paper. All authors discussed the results and commented on the manuscript.

## Data and materials availability

All data are available in the manuscript or the supplementary materials.

## SUPPLEMENTARY

Materials and Methods

Supplementary Text

Figs. S1 to S11

