## Supplementary material for "Discovery of Phenyl-β-D-glucuronide Medical Function for in Vivo Producing Handheld Gas Sensor Detectable Phenol-like Breath Marker: The Future of Induced Volatolomics in Cancer Risk Pre-warning": Supplmental Materaisl and Methods

**MATERIALS AND METHODS**

**Synthesis of** phenyl-β-D-glucuronide

*Step1: 2,3,4-Tri-O-acetyl-1-O-phenyl-β-D-glucopyranuronic acid methyl ester.*

Methyl 1,2,3,4-tetra-O-acetyl-β-D-glucuronate (10.00 g, 26.6 mmol) was dissolved in CH_2_Cl_2_ (150 mL) in an N_2_ atmosphere. Phenol (6.25 g, 66.43 mmol) and SnCl_4_ (3.46 g, 13.29 mmol) were then added, and the reaction mixture was left to stir overnight at room temperature. The mixture was then diluted with dichloromethane (150 mL), then saturated aqueous NaHCO_3_ solution (300 mL) was added and the resulting mixture was stirred for 30 min. The mixture was filtered through celite, the layers were separated, and the organic layer was dried (Na_2_SO_4_), and filtered. The solvent was then removed and the residue was purified by column chromatography (EA/PE gradient elution) to afford off-white solid, yield 4.56 g (41.8%).

^1^H NMR (400 MHz, Chloroform-*d*) δ 7.35 – 7.24 (m, 3H), 7.13 – 7.00 (m, 2H), 7.00 – 6.96 (m, 1H), 5.40 – 5.35 (m, 1H), 5.35 – 5.24 (m, 2H), 5.15 (d, *J* = 7.1 Hz, 1H), 4.23 – 4.13 (m, 1H), 3.73 (s, 3H), 2.09 – 1.99 (m, 9H). (Supplementary Figure 1a)

^13^C NMR (101 MHz, Chloroform-*d*) δ 170.13 , 169.35 , 169.24 , 166.89 , 156.72 , 129.63 , 123.54 , 117.09 , 99.23 , 77.23 , 72.70 , 71.90 , 71.10 , 69.15 , 52.98 , 20.63 , 20.53 . (Supplementary Figure 1b)

MS: 428.1 [M+NH_4_]^+^. (Supplementary Figure 2a)

IR (neat, cm^-1^): 2951, 1759, 1597, 1492, 1438, 1381, 1296, 1222, 1080, 1045, 914, 767, 694. (Supplementary Figure 2b)

*Step2: Phenyl-β-D-Glucopyranosiduronic Acid.*

2,3,4-Tri-O-acetyl-1-O-phenyl-β-D-glucopyranuronic acid methyl ester (4.50 g, 11.0 mmole) was dissolved in methanol (45 mL) and NaOMe (0.24 g, 4.4 mmol) was added. After 2 h Amberlite was added, the reaction mixture was filtered and the solvent removed and the residue was purified by pre. HPLC to afford white solid, yield 1.04 g (35.1%).

^13^C NMR (101 MHz, D_2_O) δ 174.19 , 156.54 , 129.96 , 123.45 , 116.71 , 100.15 , 75.57 , 75.24 , 72.72 , 71.54 . (Supplementary Figure 3a)

MS: 288.1 [M+NH_4_]^+^. (Supplementary Figure 3b)

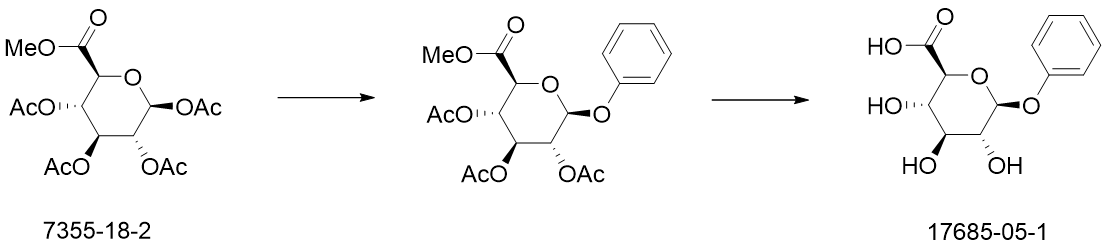

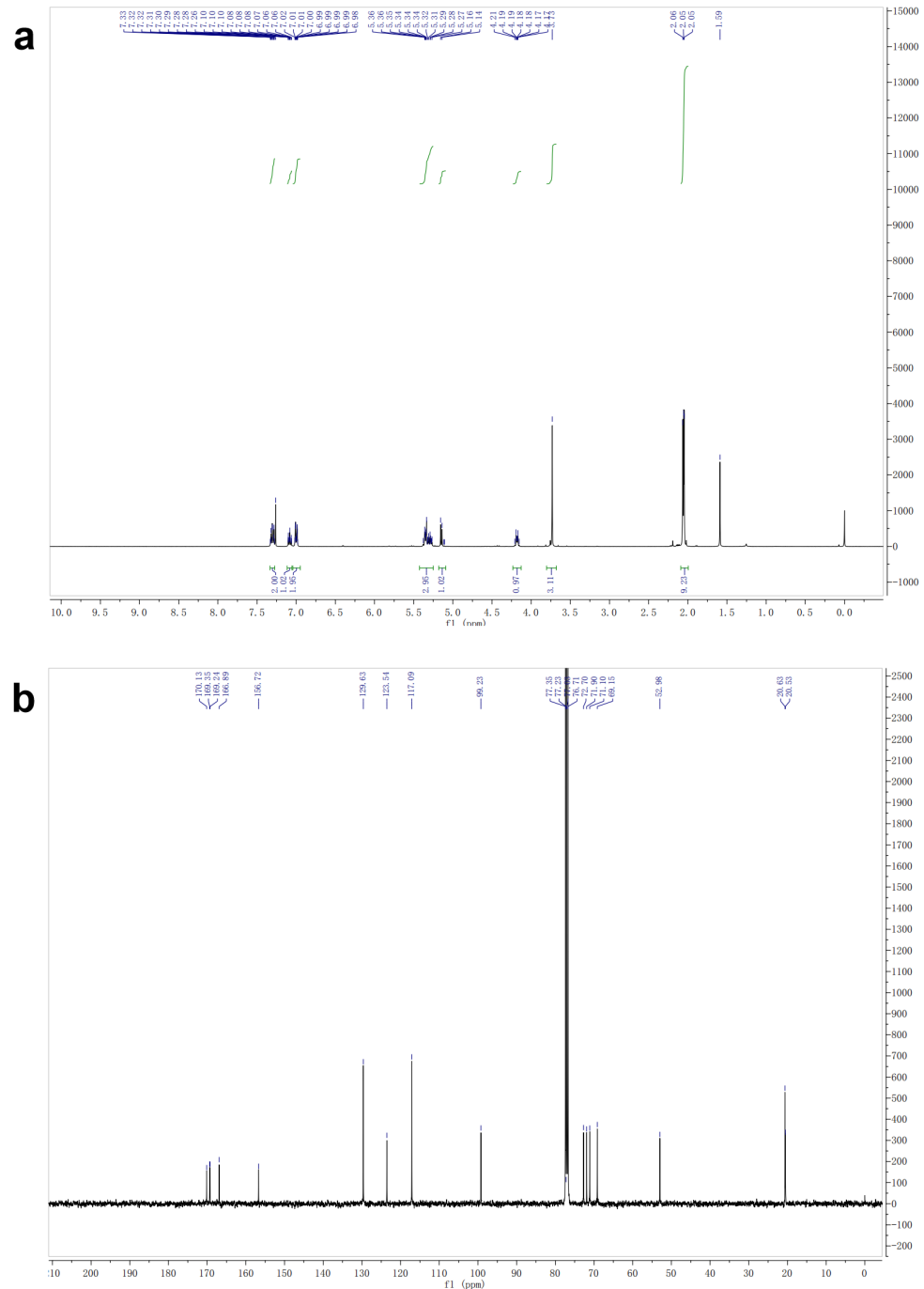

Supplementary Figure 1. NMR characterization of intermediate. a. HNMR and b. CNMR spectra of the intermediate

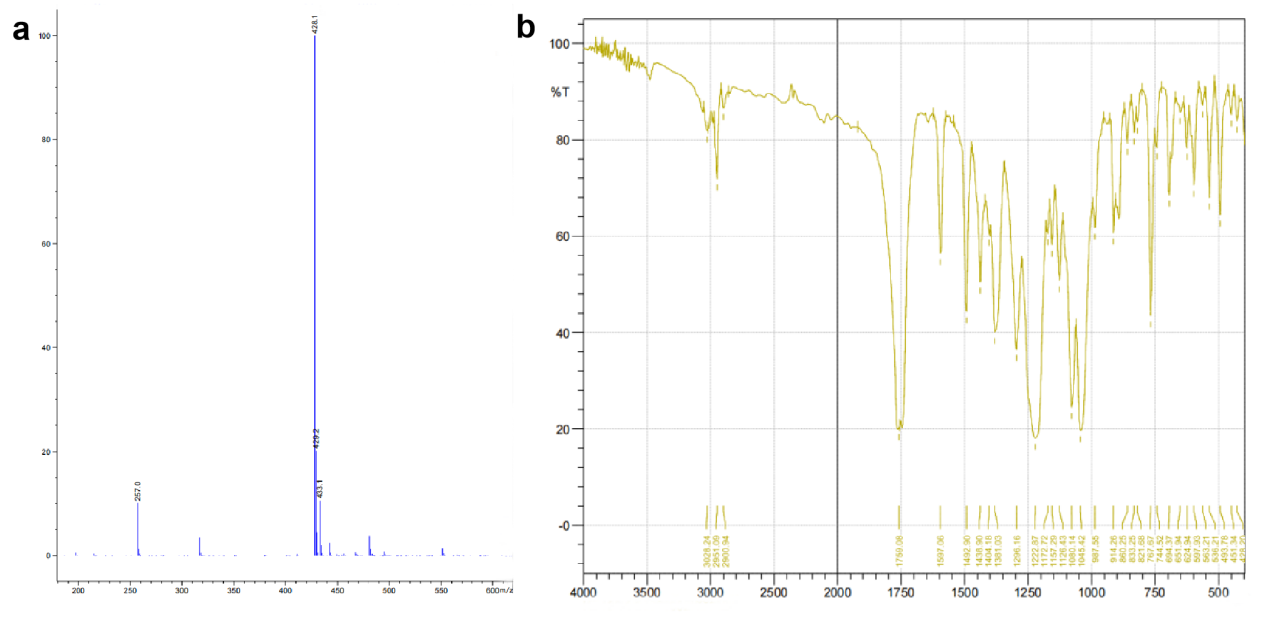

Supplementary Figure 2. Molecular structure and functional group analysis. a. MS and b. IR spectra of the intermediate

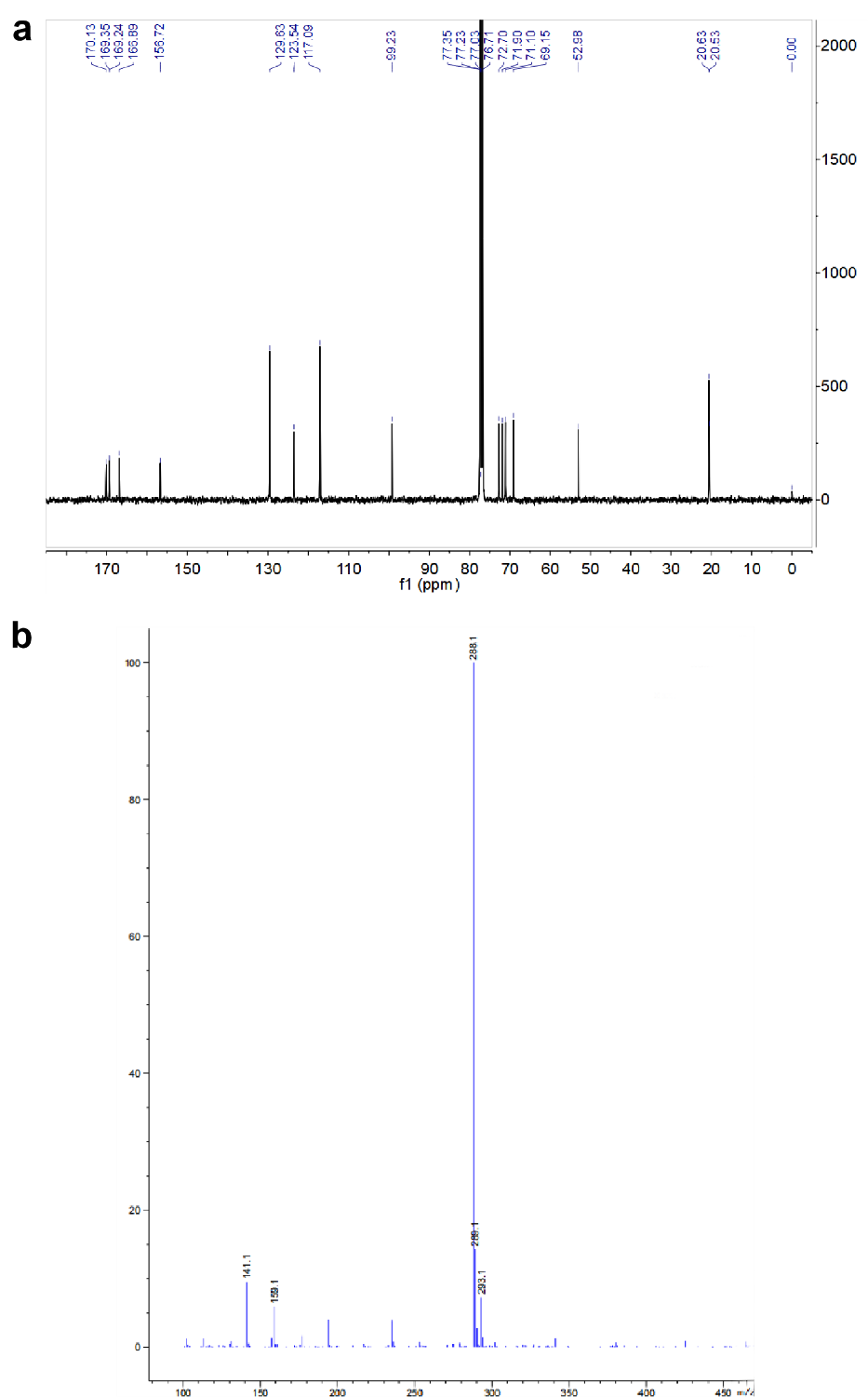

Supplementary Figure 3. Molecular structure analysis. a. CNMR and b. IR spectra of the synthesized phenyl-β-D-glucuronide product

**In vitro enzyme-catalyzed reaction products investigation**

(1) Solution preparing: The buffer solution was prepared using sterile water, with the main components of 75 mM potassium dihydrogen phosphate (KH2PO4) and 1% bovine serum albumin (BSA), and adjust the pH to 6.8 using 1 M potassium hydroxide (KOH). Prepare solutions of 4-nitrophenyl-β-D-glucoside and phenyl-β-D-glucoside separately using deionized water, each with a concentration of 0.75 g/L. (2) Enzyme-catalyzed reaction: The enzyme-catalyzed reaction was performed in two 20 mL glass sample bottles and sequentially added with 2.6 mL of deionized water, 2 mL of buffer solution, and 1 mL of 4-nitrophenyl-β-D-glucoside or phenyl-β-D-glucoside as the substrate solution. After thorough mixing, the mixture was stabilized in a 37°C water bath for 10 minutes. 2 μL of glycerol solution containing β-glucosidase (enzyme content: 500 U) was added into one bottle as the experiment reaction system while 2 μL of glycerol to the control reaction system, and quickly tighten the lid and react in a 37°C water bath for 1 hour. (3) Enrichment and analysis of gas molecules: The pre-activated solid-phase microextraction (SPME) was used to enrich the gas molecules at room temperature for 30 minutes, the reaction solution was then analyzed with high-performance liquid chromatography-mass spectrometry (LCMS). Gas components are analyzed using gas chromatography-mass spectrometry (GCMS), with the extraction core desorbed at 260°C for 5 minutes at the injection port. The chromatographic column was DB-WAX (30 m × 0.25 μm × 0.25 mm), with an initial temperature of 40°C held for 5 minutes, followed by a ramp-up at 10°C/min to 250°C and a further 5-minute hold. Mass spectrometry is conducted with a full-range scan from 29 to 400 amu using high-purity helium as the carrier gas at a 1 mL/min flow rate. Substances detected are analyzed using the NIST14 spectral library.

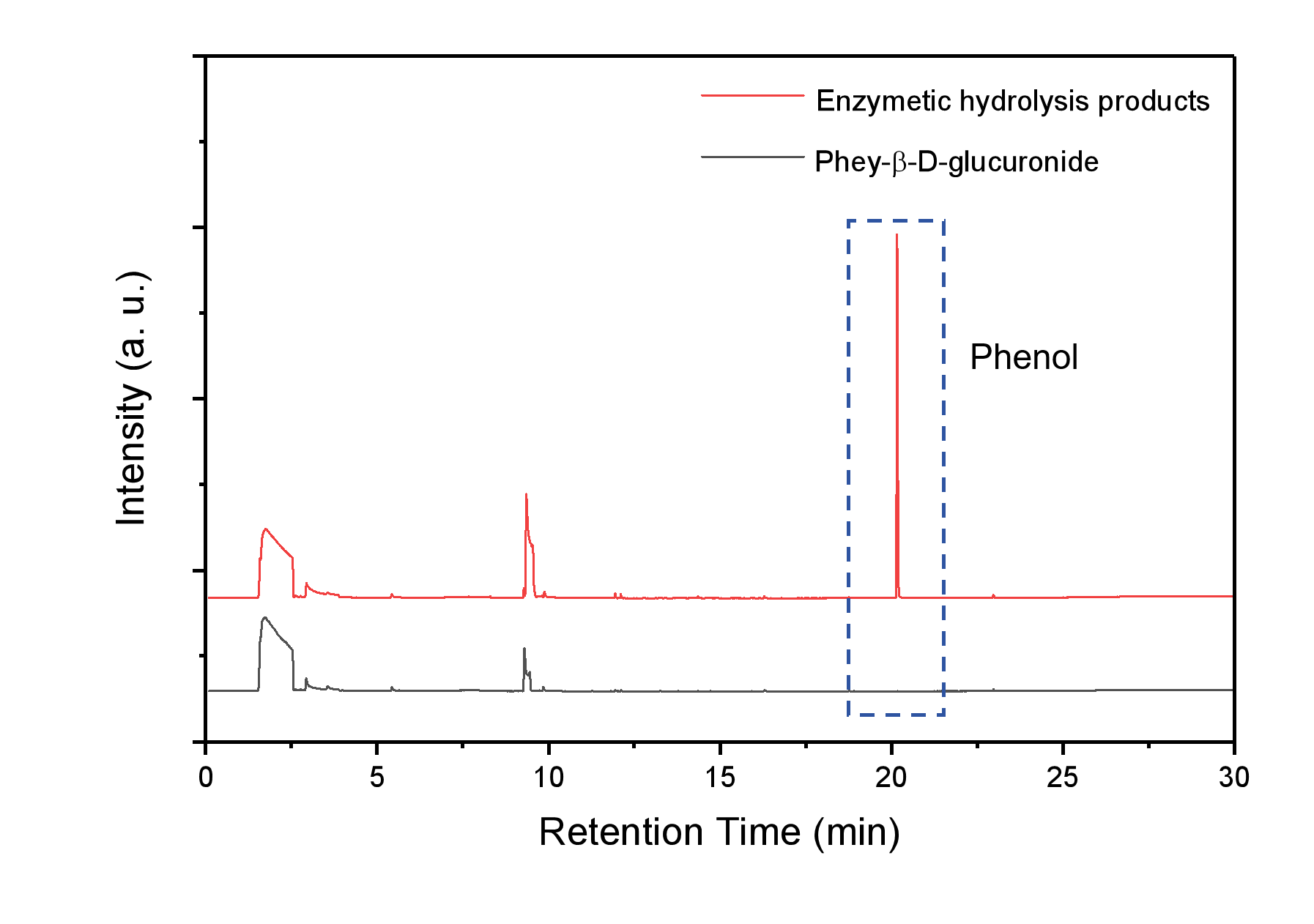
Supplementary Figure 4. Verification of phenol generation during enzymatic reaction. Upper gas phase sample is directly analyzed by LC to confirm the generation of phenol vapor.

**Toxicity investigation of** phenyl-β-D-glucuronide like exogenous agent

The cytotoxicity of drugs was characterized with the Cell Counting Kit-8 (CCK-8) assay. The complete culture medium for cultivating gastric mucosal epithelial cells included a high-glucose DMEM culture medium, 10% fetal bovine serum, and 1% penicillin-streptomycin (double antibiotics). Gastric mucosal epithelial cells were resuscitated in 15 mL of complete culture medium, placed in a 37 ^o^C constant temperature incubator with 5% carbon dioxide, and the culture medium was changed every 1-2 days. Cells were seeded in a 96-well plate at a concentration of 5*10^4^ cells/mL, with 100 μL per well, and 5 replicate wells were set for each sample group. After 30 h of cell culture, the culture medium was discarded, and the DMEM culture medium containing phenyl-β-D-glucoside was added. The drug concentrations were set as 0 μg/mL, 0.1 μg/mL, 1 μg/mL, 10 μg/mL, 100 μg/mL, 500 μg/mL, and culture medium without seeded cells, with each well having 100 μL of culture medium. After a total incubation of 40 h, the culture medium was discarded, and 100 μL of DMEM culture medium containing 10% CCK-8 was added to each well. After incubating for 2 h, the absorbance values (OD) at 450 nm wavelength for each well were measured using an enzyme marker, and cell viability was calculated based on the absorbance values. Additionally, mice subjected to the inductive exhalation analysis were euthanized, and major organs were sectioned for Hematoxylin and Eosin (HE) staining to conduct histopathological examination, characterizing whether repeated drug injections had toxic side effects on the organs. The formula for calculating cell viability is:

$$Cell Viability=\frac{OD_{Drug}-OD_{Blank}}{OD_{Control}-OD_{Blank}}*100\%$$

**Cell culture and** **tumor model construction**

Cell culture: The human gastric cancer cell line (MGC-803) was obtained from the Cell Bank of the Chinese Academy of Sciences. Cells were cultured in Dulbecco's Modified Eagle Medium (DMEM) supplemented with 10% fetal bovine serum and 1% penicillin-streptomycin (double antibiotics) in a CO_2_ incubator at 37°C with 5% CO_2_.

Mice breeding: Six-week-old female BALB/c-Nude nude mice were purchased from Shanghai JSJ Experimental Animals Co., Ltd., with an SPF level, and an initial weight of 17±1.5 g at the time of purchase. The mice were housed in clean-grade animal rooms in individual cages, with ventilation maintained through a central control system. The indoor temperature was kept at 25±2 ^o^C, and both food and water were provided after high-pressure sterilization. After one week of acclimatization, during which the mice reached a weight of 20±2 g, their growth was observed daily. The mice were randomly divided into two groups: the cancer group and the healthy group. The cancer group was used to establish a subcutaneous gastric cancer model with MGC-803 cells.

Tumor Model Construction: MGC-803 cells were revived in 15 mL complete culture medium and placed in a constant temperature incubator at 37 °C with 5 % carbon dioxide, with the culture medium changed every 1 to 2 days. The adherent gastric cancer cells were digested with trypsin, centrifuged, collected, resuspended, and counted. The cells were then dispersed in a 15 mL complete culture medium at a density of 1*10^6^ cells/mL in cell culture bottles. After reaching confluence, the cells were digested with trypsin, centrifuged, washed twice with PBS to remove residual serum, and resuspended in PBS with a density of approximately 5*10^7^ cells/mL. Each mouse was inoculated with 100 μL of cell suspension. The cell inoculation process was completed within half an hour, during which the cells were placed on ice to reduce metabolic activity and maintain cell viability. Tumors were implanted in the well-vascularized area in the posterior part of the armpit, and the day of tumor implantation was recorded as Day 0. After tumor formation, the tumor size and mouse weight were measured every 3 to 4 days using calipers and electronic weigher, measuring both the longest and shortest dimensions of the tumor. Tumor volume was calculated using the following formula:

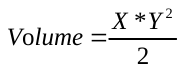

Here, X represents the longest dimension of the tumor, and Y represents the shortest dimension of the tumor, measured in millimeters. According to animal ethics standards, when the tumor volume of mice exceeds 2 cm^3^ or if necrotic tissue develops in the tumor site, it may adversely affect the normal life of the mice, and euthanasia should be considered for the well-being of the mice.

**Breath sample collection**

Mouse exhaled breath was analyzed for gas composition using GCMS, and rapid tumor screening was achieved through signal detection using self-developed electrochemical sensors. For this project, two types of exhaled breath collection devices were designed based on experimental requirements: an exhaled breath extraction device and an exhaled breath collection device.

(1) Mouse exhaled breath extraction device

The exhaled breath extraction device enriches volatile organic compounds (VOCs) from mouse exhaled breath onto an SPME fiber for the analysis of exhaled breath composition and relative content before and after mouse administration. The device consists of a large glass container with a volume of approximately 400 mL. The bottleneck was sandblasted, and the device was sealed with a sealing gasket, silicone plug, and fixing clip to ensure airtightness. The device was first cleaned and dried before use. Cleanliness and airtightness tests are conducted initially. In the cleanliness test, high-purity nitrogen gas was introduced into the device, sealed for 1 hour, and then the gas adsorbed on the SPME fiber was analyzed by GCMS. The composition of the adsorbed substances was compared with a blank SPME fiber to confirm that the device itself does not generate impurity gases. In the airtightness test, the sealed device was injected with gas, and performed a soap bubble test. No bubbles are produced at any part of the device, and the piston of the syringe returns to its original position after release, indicating good sealing performance of the device.

Place the mice that need to undergo exhaled breath component extraction into a glass container. The entire apparatus is placed in a constant temperature environment at 25°C and sealed for 1 hour, with oxygen supplementation every 15 minutes. Using a glass syringe through a silicone plug, inject 10 mL of pure oxygen (99.9%). After 1 hour of sealing, insert a 75 μm Car/PDMS extraction fiber into the glass container through the silicone plug, exposing the black extraction fiber of SPME. The extraction was carried out for 30 minutes in a constant temperature environment at 25°C, and oxygen supplementation was performed periodically following the above process.

(2) Mouse exhaled breath collection device

The mouse exhaled breath collection device was used to collect the mouse's exhaled breath into a gas bag, facilitating sensor testing experiments. In sensor experiments, the exhaled breath must pass through the sensor's sensitive electrode. Therefore, the mouse's exhaled breath needs to be collected into a gas bag for sensor testing. To meet the requirements of this experiment, a gas collection device is assembled, including a 500 mL sandblasted glass syringe, silicone tubing, and a Tedlar gas bag. The glass syringe was used to collect the mouse's exhaled breath, and it was thoroughly cleaned and dried to remove any potential impurities and sealed with silicone tubing, making it convenient to supplement oxygen when collecting breath from the mouse. The Tedlar gas bag, used for collecting and storing gas, was thoroughly dried and washed three times with high-purity nitrogen gas to remove impurities adsorbed on the inner wall of the bag.

In the gas collection system, both the oxygen bag and the sampling bag were connected to the syringe through a silicone tube, and the gas volume was determined by the syringe's scale. The overall cleanliness and sealing are tested. For cleanliness testing, high-purity nitrogen gas is introduced into the glass syringe and the sampling bag. After sealing for 1 hour, the gas from both is separately adsorbed using an extraction fiber. The gas is then analyzed by GCMS through headspace injection, and the results are compared with the analysis of a blank extraction fiber. In the sealing test, 500 mL of gas was injected into the syringe, and the silicone tubing in the device was clamped. The gas collection syringe is placed vertically for 1 hour. The scale position of the piston remains at 500 mL, confirming the good sealing performance of the device.

Place the targeted mouse into the glass syringe, push the piston to the 450 mL mark, connect the other end of the silicone tubing attached to the gas outlet to the oxygen bag, and seal with a clamp. The entire apparatus was placed in a 25°C environment, and every 15 minutes, the sealed clamp and the gas bag switch, and move the piston to supplement 10 mL of oxygen. After 1 hour of sealing, remove the oxygen bag, connect an empty sampling bag, and slowly push the gas from the syringe into the sampling bag by pushing the piston. Ensure sufficient space for the mouse during this process. After the gas collection, remove the mouse and place it in a regular mouse cage immediately.

**Analysis of exhaled breath components**

The mouse's exhaled breath was extracted and analyzed with SPME and GCMS to analyze the changes in mouse exhaled breath composition before and after induction drug treatment. The SPME extraction fiber needs to undergo conditioning before use to remove impurities adsorbed during storage, with a conditioning program set at 260°C for 30 minutes, helium carrier gas flow rate at 1 mL/min, and this conditioning process conducted in the injection port of the GC-MS instrument. After the extraction fiber conditioning was complete, the column temperature was set to 245°C for an additional 30 minutes. The testing method was as described earlier, with determination criteria for induced characteristic biomarkers having a matching probability no less than 90%, and a retention time difference with standard substances within 0.1 minute. Analysis was performed on characteristic biomarkers before and after drug administration, as well as other substances in exhaled breath that show significant changes.

**Fabrication and characterization of YSZ-based electrochemical phenol sensor**

(1) Comparison of the gas adsorption capability for various metallic oxides

The comparison of adsorption capability for the various metallic oxides (Fe_2_O_3_, CeO_2_, In_2_O_3_, SnO_2_, TiO_2_, Cr_2_O_3_, WO_3_, Co_3_O_4_, ZnO, NiO) to each studied gas is implemented by a analyzer of interface thermodynamic/kinetic parameters (ITKP, Xiamen High-End MEMS Technology Co. Ltd., China).

(2) Sensor fabrication

The sensor was fabricated as a three-electrode electrochemical sensor based on yttria-stabilized zirconia (YSZ). It has a flat plate structure, with the sensing electrode (SE), reference electrode (RE), and counter electrode (CE) all located on the same side of the plate, connecting to one end of printed silver wires. On the other side of the flat sensor was an alumina heating element, which is intimately bonded to the YSZ solid electrolyte through co-firing. The printed silver wires on this side are connected to the heating power source. The printed silver wires on both sides of the sensor accurately correspond to a customized fixture, which was used to transfer signals to the signal detection device and provide heating voltage. In this project, chromium oxide was used as the sensing electrode, manganese dioxide as the reference electrode, and platinum slurry as the counter electrode. The overall dimensions of the sensor is 34*4*1 mm³.

The brief sensor fabrication processes are as follows:

1) Binder preparation: Mix turpentine alcohol and ethyl cellulose in a ratio of 94:6 by mass to prepare a turpentine alcohol slurry.

2) Preparation of reference electrode: Grind a mixture of turpentine alcohol slurry and manganese dioxide powder in a mass ratio of 1.2:1 using an agate mortar to obtain a reference electrode slurry. Coat the reference electrode slurry onto the reference electrode lead based on yttria-stabilized zirconia.

3) Preparation of counter electrode: Coat platinum slurry onto the counter electrode wire.

4) Sintering of reference and counter electrodes: The devices were first dried at 130°C for 4 hours and then placed in a muffle furnace for sintering. The sintering temperature profile was as follows: ramp up from room temperature to 1400°C at a rate of 3°C/min, hold at 1400°C for 2 hours, and then naturally cool to room temperature after the sintering process.

5) Preparation of the sensing electrode: Grind a mixture of turpentine alcohol slurry and chromium oxide powder in a mass ratio of 1:1 using an agate mortar to obtain a sensing electrode slurry. Coat the sensing electrode slurry onto the lead of the sensing electrode.

6) Sintering of the sensing electrode: The devices were first dried at 130°C for 4 hours and placed in a muffle furnace for sintering. The sintering temperature profile is as follows: ramp up from room temperature to 1050°C at a rate of 3°C/min, hold at 1050°C for 2 hours, and then naturally cool to room temperature after the sintering process.

(3) Details of gas blending

The testing system mainly consists of the sensor, quartz testing chamber, digital multimeter, power supply, gas cylinders, and a gas distribution system. The gas distribution system has nine interfaces, which can be connected to nine gas cylinders (replaceable) through pressure-reducing valves. The gas switches and flow direction were controlled by two-way/three-way valves in the pipeline. The gas flow rate was controlled by seven mass flow controllers (MFC) arranged in the pipeline. The real-time flow was adjusted and monitored by the Muti-Digital MFC software on the computer. The digital multimeter was connected to the computer through an I/O interface transmission line, and the Agilent signal acquisition software was used to collect and display the response curve of the sensor in real time. The entire system can dynamically configure gases of different concentrations and compositions, exposing the sensor to the atmosphere of the test gas to achieve real-time collection of response signals.

To align with real-world application scenarios, the sensor was exposed to a simulated air environment, and the response signal in this environment was considered as the baseline state for the sensor. The main principle of sensor testing relies on the redox reactions at the interface, directly correlated with the concentration of oxygen. In air, the primary factor affecting the sensor response signal is the concentration of oxygen. Therefore, the simulated air contains 21% oxygen, with the rest being nitrogen. Additionally, when diluting samples of different concentrations or creating gas mixtures, it is essential to ensure that the final oxygen concentration in the gas is 21%. For dilution, three gases are used: the original test gas, 50% oxygen, and air. The total gas flow rate was maintained at 100 sccm, and the proportions of different gases in the mixture were controlled by the gas flow rates. This was achieved through mass flow controllers to adjust the gas concentration, with the calculation formula as follows:

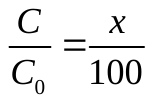

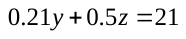

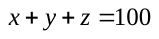

Where C_0_ is the original concentration of the test gas, C is the target concentration of the test gas; x, y, and z represent the actual gas flow rates of the test gas, air, and oxygen, respectively. Based on the above principal formula, the numerical values of the flow rates for the three required gases during gas dilution can be obtained as follows:

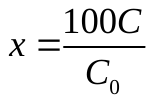

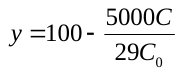

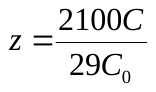

The original gases used in this experiment are listed in Supplementary Table 1.

**Supplementary Table 1.** Original gases used for preparing sample gas

| **VOCs** | **CAS** | **Gas Concentration** | **Manufacturer** |
| --- | --- | --- | --- |
| Phenol | 108-95-2 | 1.2 ppm | Shanghai Wei Chuang Standard Gas Analysis Technology Co., Ltd. |
| Oxygen | 7782-44-7 | 50 % |  |
| Air | N/A | N/A |  |
| Acetone | 67-64-1 | 20.3 ppm |  |
| Styrene | 100-42-5 | 15.0 ppm |  |
| Hexanal | 66-25-1 | 18.5 ppm |  |
| Ethanol | 64-17-5 | 18.7 ppm |  |
| n-Butanol | 71-36-3 | 20.1 ppm |  |
| Ethylbenzene | 100-41-4 | 16.7ppm |  |
| 2-Pentanone | 107-87-9 | 20.1 ppm |  |

Note: High-purity nitrogen gas is used as the balance gas for all gases

(3) Sensor characterization

Both SEs and Mn-based RE of the sensor were simultaneously exposed to the base gas (diluted with air base) or the sample gas containing each of various VOCs to evaluate the gas sensing characteristics. Since the main research objective of the this study is to develop a phenol sensor for the application of breath analysis, acetone, styrene, hexanal, ethanol, n-butanol, ethylbenzene which are widely reported to be found in animal breath are selected as the interference gases. The base gas uses a volume ratio of 21%O_2_ mixed with N_2_ standard gas, maintaining a gas flow rate of 100 sccm (MFC: CS200 SCCM, Beijing Qixing Huachuang Electronics CO., Ltd). Three standard gases with a volume ratio of 50% O_2_ mixed with residual N_2_, phenol (or acetone, styrene, hexanal, ethanol, n-butanol, ethylbenzene) mixed with residual N_2_ and 21 % O_2_ mixed with residual N_2_ (Shanghai Weichuang Standard Gas Analysis Technology CO., Ltd.) were selected by using a mass flow controller (MFC: CS200SCCM, CS100SCCM, CS100SCCM, Beijing Qixing Huachuang Electronics CO., Ltd.) control the volume ratio of O_2_ to maintain at 21 % and a total gas flow rate of 100sccm. The electric potential difference (ΔV, ΔV = V_sample_ _gas_ − V_base gas_) between SE and RE is recorded by using an electrometer (34970A, Agilent, USA). The operating temperature ranges from 400 to 500 °C. The carrier gas's background relative humidity (RH) was controlled by precisely blending dry air with air that was fully saturated with moisture (RH 100%). The hygrometer (4185 Traceable, USA) was used to monitor the ratio between the partial pressure of water vapor and the equilibrium vapor pressure of water in the mixture. The findings indicate that under steady environmental conditions within the range of 25-27°C, the relative humidity test yields a result of around 70%. All subsequent sensing test environments are conducted under the same temperature and humidity conditions.

**Pre-warning of cancer risk with the handheld breath-analyzer**

Air bag that filled with the breath samples derived from healthy or tumor bearing mice was directly connected to the inlet of the glass rotor flow-meter, and the gas flow rate was set to 45 mL/min. The outlet of the flow-meter was connected to the air pump, which was then connected to the handheld breath-analyzer.

Phenyl-β-D-glucuronide solution is prepared using PBS buffer with a 400 μg/mL concentration. The healthy group and tumor group mice were intravenously injected with the drug, with an induction injection volume of 200 μg/kg. Each mouse was injected with approximately 100 to 120 μL of the phenyl-β-D-glucuronidesolution based on its weight. During the testing process, the surrounding air was directly sampled as background gas. After the baseline stabilizing, the airbag containing mouse exhalation was connected, and a 5-minute timer was started. The response signal of the sensor to the sample gas was obtained by subtracting the baseline value. After testing the sample gas for 5 minutes, the airbag containing the test air was removed. The process was repeated after a 5-minute interval. Before and after phenyl-β-D-glucuronide injection, mouse exhalation was collected using the mouse exhalation collection device, and the sensor was used to test its response signal. The change in the response signal before and after administration was calculated, and each mouse's exhalation sample was generally tested in two cycles.

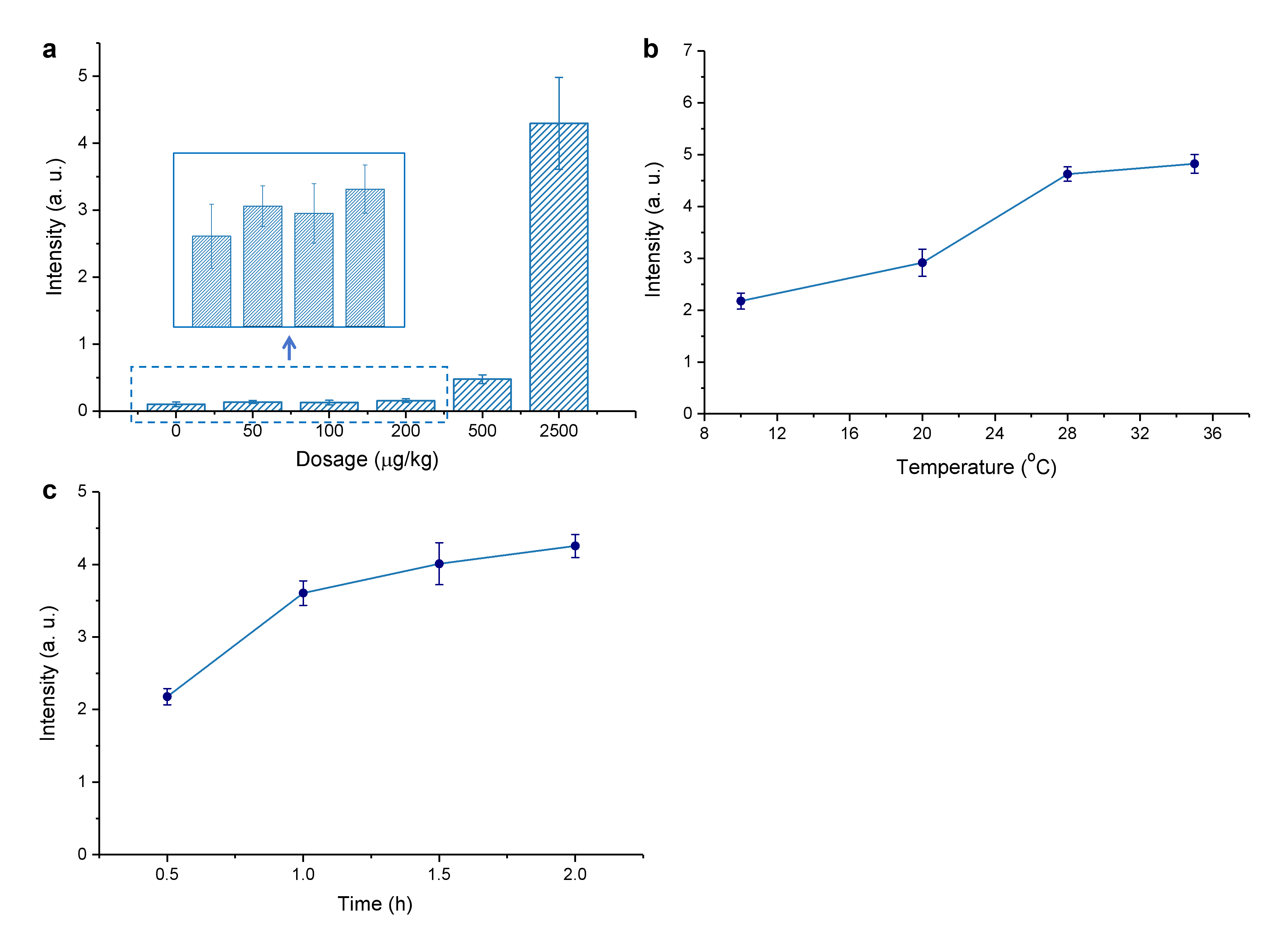

Supplementary Figure 5. Experimental parameters optimizing. a, Variation of the induced phenol level in breath sample derived from health mice (n = 5 for each administration) at the phenyl-β-D-glucuronide dosage in the range of 0-2600 μg/kg. A minimal background signal of phenol in exhaled breath of healthy animals is found at the injection of a very low 200 μg/kg of phenyl-β-D-glucuronide, suggesting the optimal dosage is around 200 μg/kg. b, Impact of the temperature for mouse incubator during sampling on phenol level in exhaled sample derived from gastric tumor bearing mice (n = 5) that dosed for 200 μg/kg of phenyl-β-D-glucuronide. c, Dependence of the induced phenol concentration in exhaled breath on the sampling time, after administrating 200 μg/kg exogenous agent at the incubator temperature of 28^o^C (n = 5).

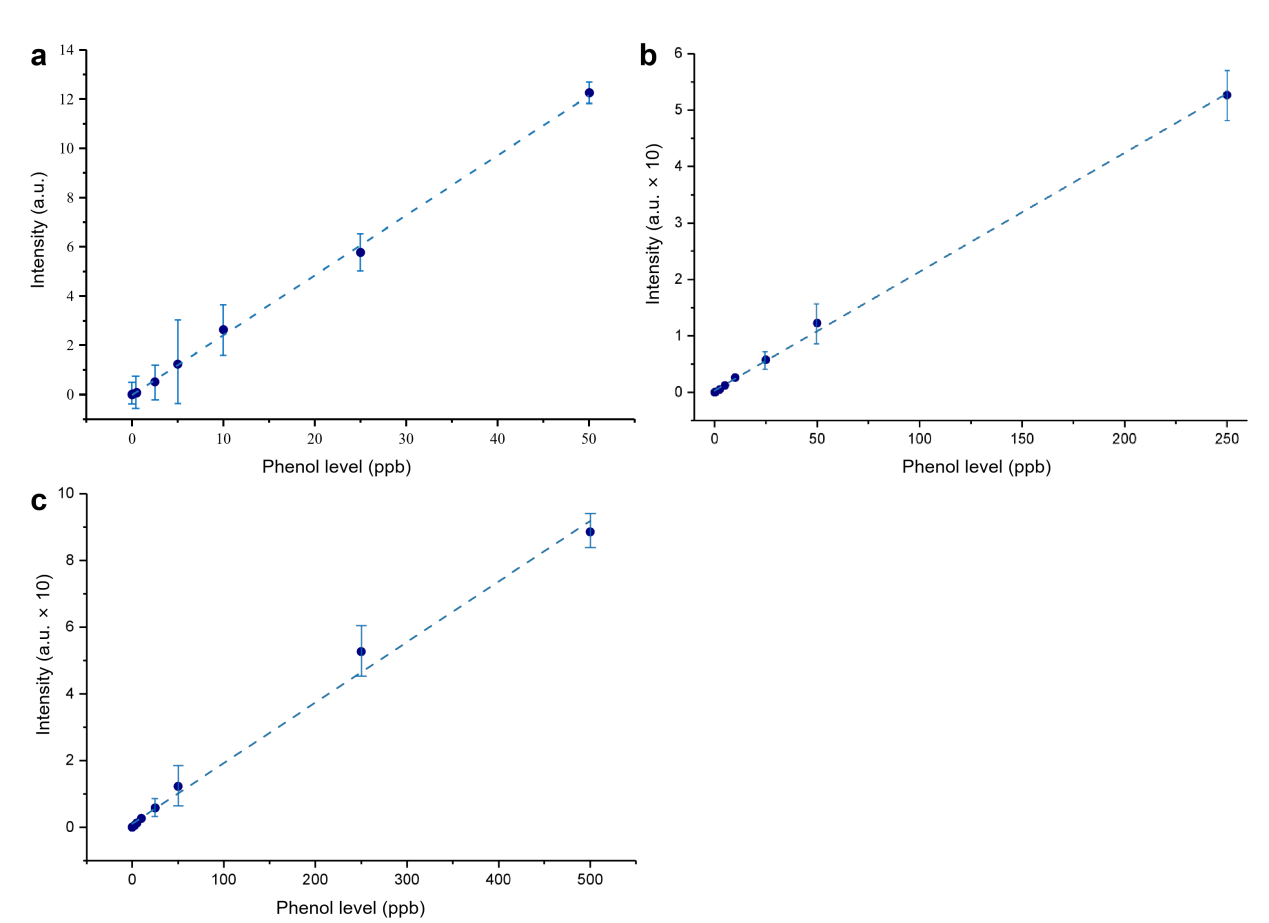
**Supplementary Figure 6. Calibration of phenol level.** Dependence of peak intensity on phenol level in the range of **a**, 0-50 ppb. **b**, **0-250 ppb. c**, 0-500 ppb.

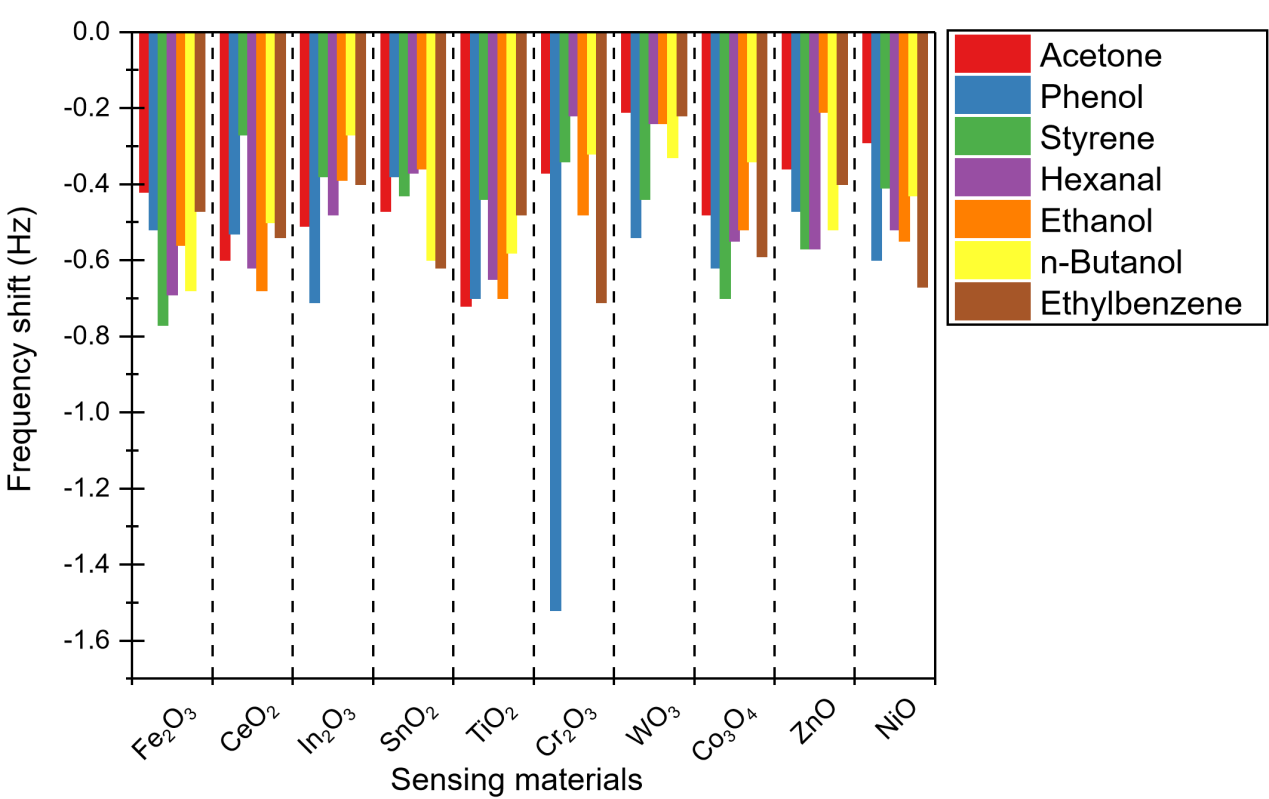

Supplementary Figure 7. Comparison of the gas adsorption capacity for various metallic oxide-based sensing materials towards acetone, phenol, styrene, hexanal, ethanol, n-hexane and ethylbenzene

Supplementary Figure 8.
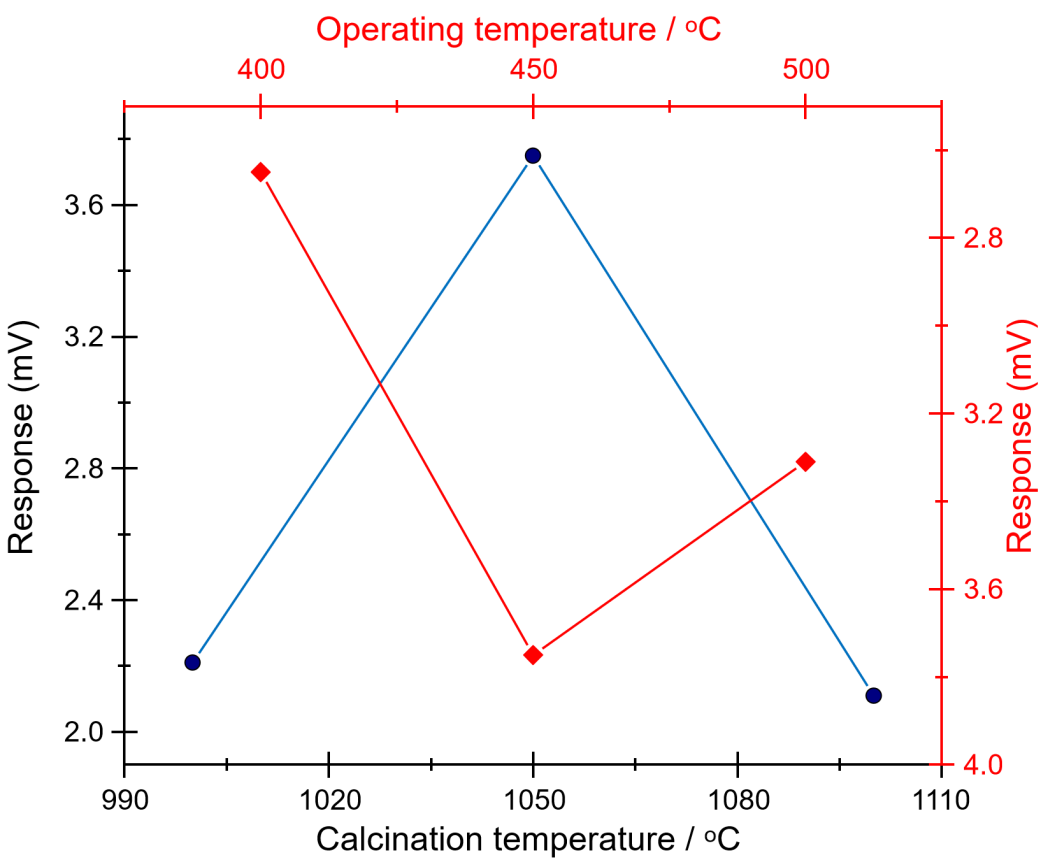
Impact of calcination and operating temperature on sensing characteristics of the YSZ-based phenol sensor utilizing Cr2O3-SE and Mn-based RE

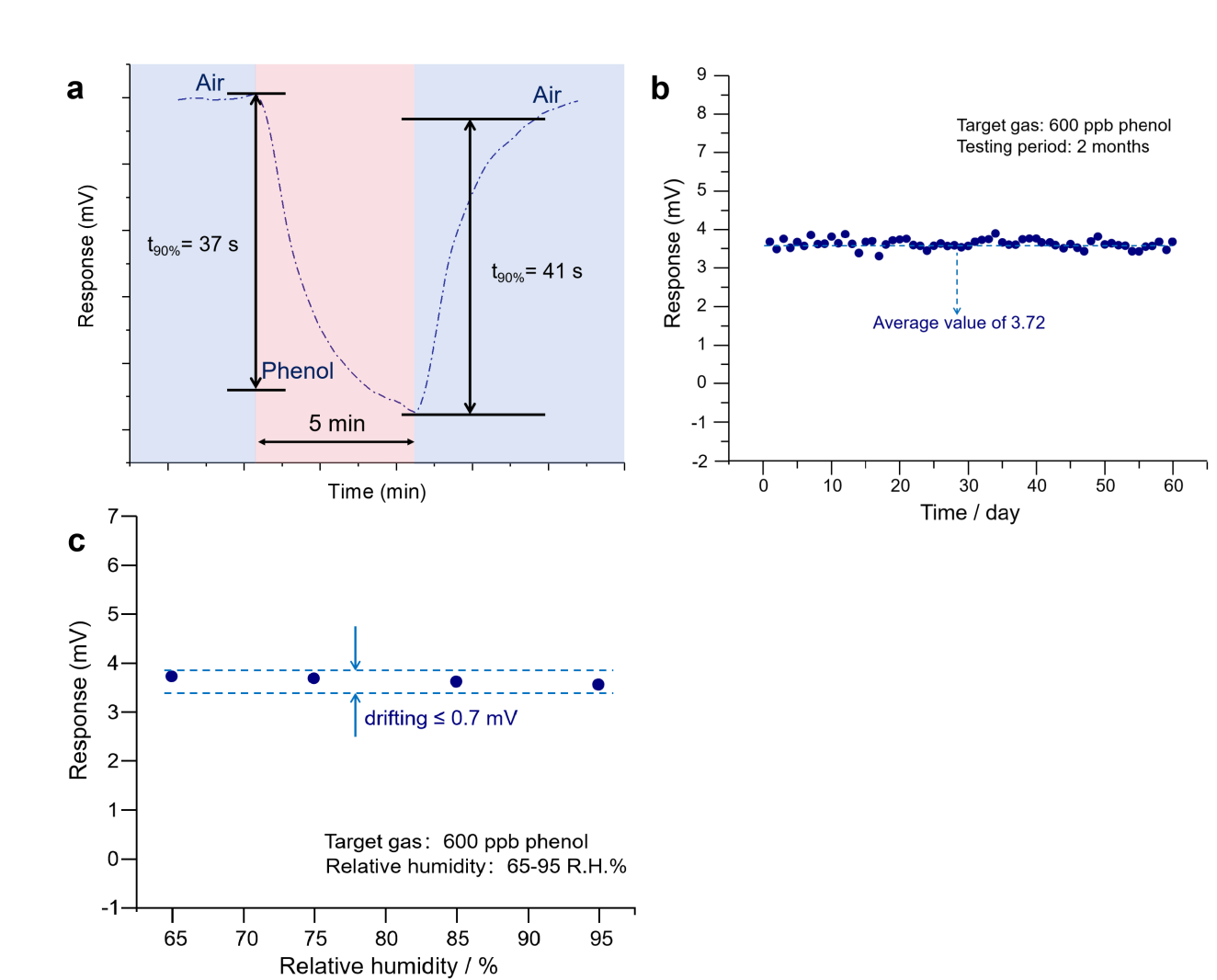

Supplementary Figure 9. **Sensing properties of the YSZ-based phenol sensor comprised of Cr_2_O_3_-SE and Mn-based RE.** **a**, 90% response/recovery time of the developed sensor. b, Long term stability of the sensor. **c**, Impact of ambient humidity on response behavior of the sensor.

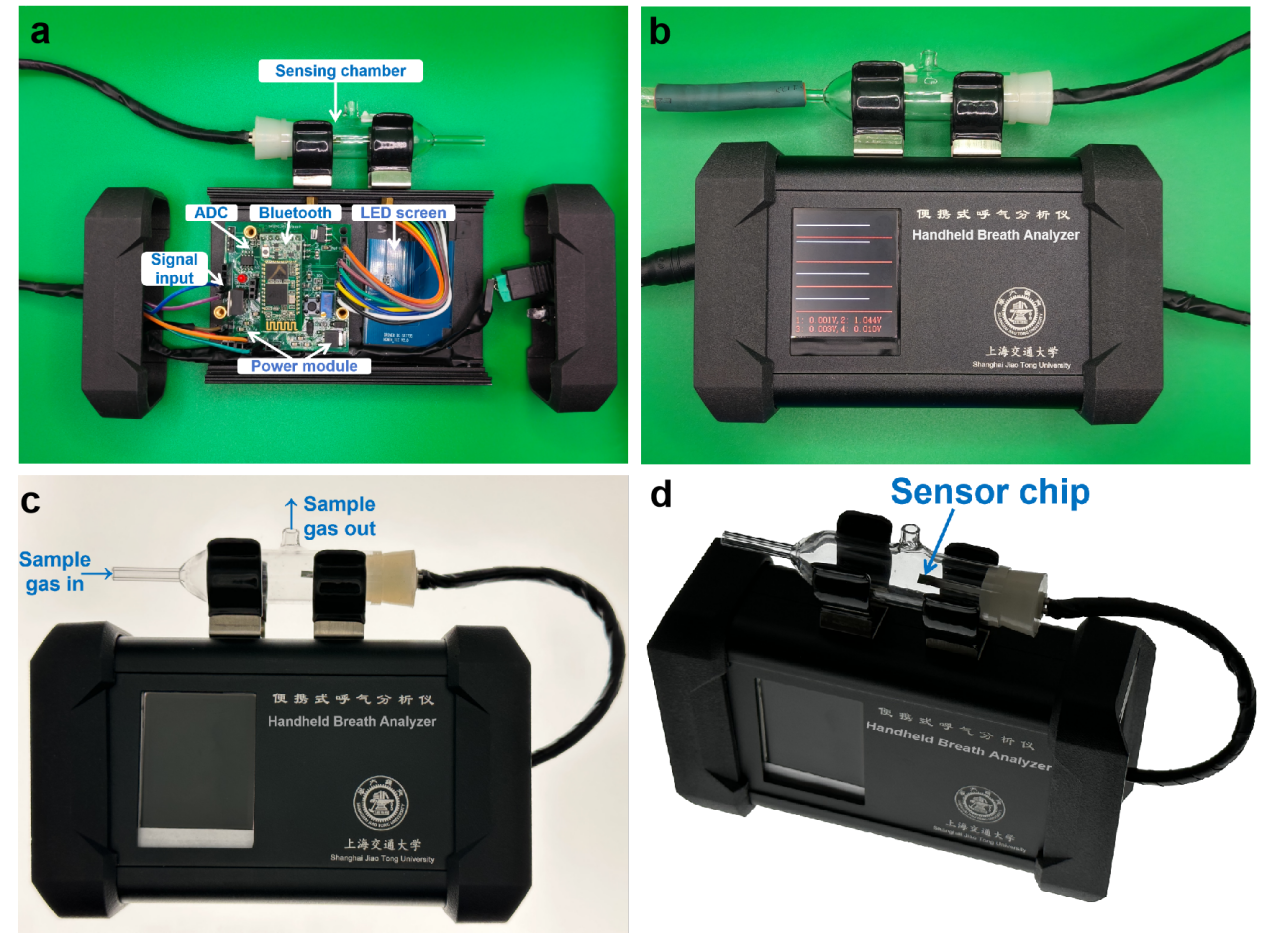
Supplementary Figure 10. Photographic images of the handheld breath-analyzer. a. Internal structure of the prototype, including display module, signal acquisition module, data storage module, sensor driver module, power supply module and wireless signal transmission module. **b.** breath-analyzer at operation mode. **c.** Sample gas transmission path in breath-analyzer. **d.** Location of the electrochemical phenol sensor in the prototype.

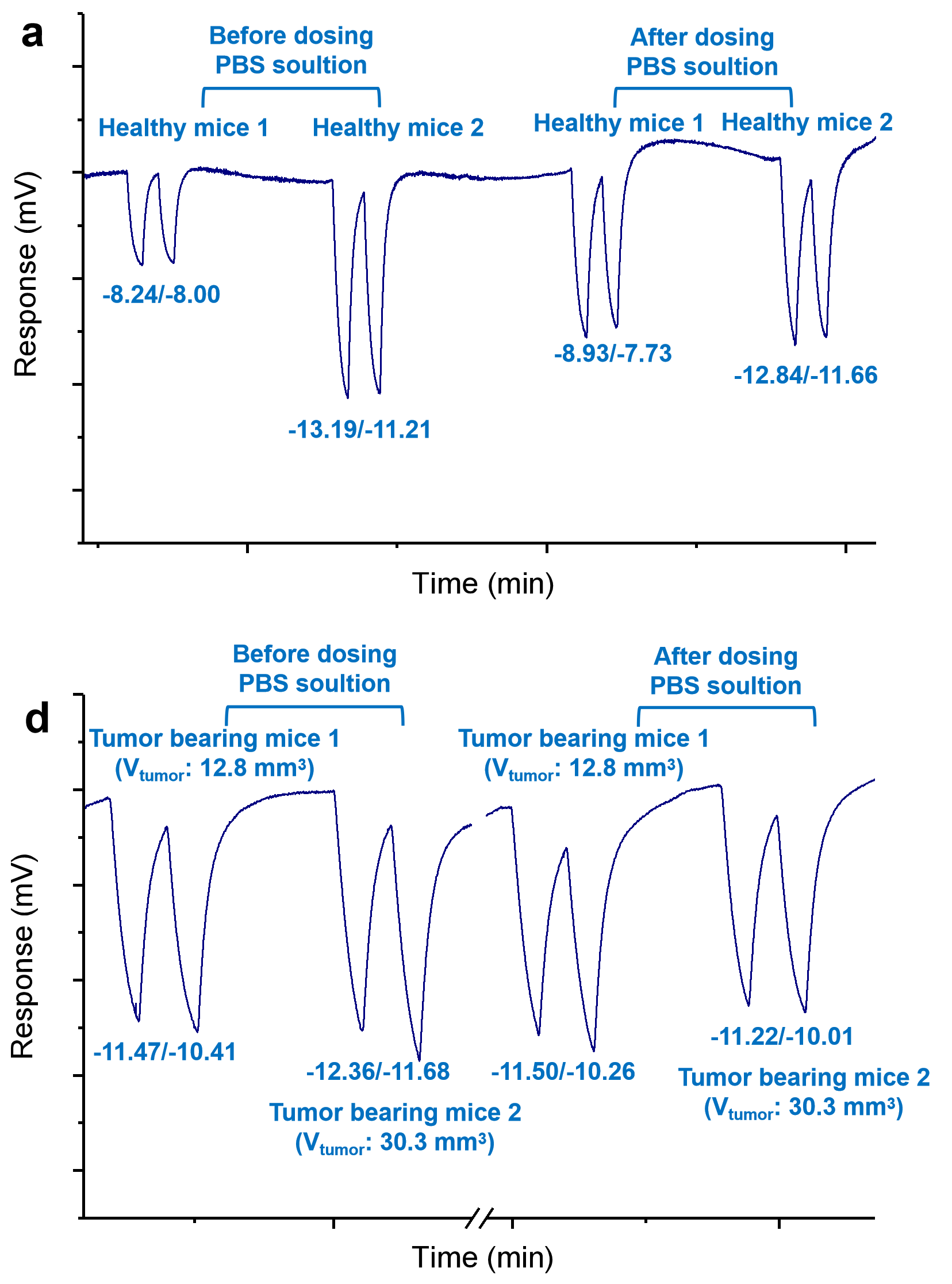
Supplementary Figure 11. **Impact of administrating PBS solution on breath signal.** Variation of the breath signal for the breath sample derived from **a**, healthy or **b**, tumor-bearing mice, after tail injecting PBS solution
