## Supplementary reports for acute systemic toxicity test. for "Discovery of Phenyl-β-D-glucuronide Medical Function for in Vivo Producing Handheld Gas Sensor Detectable Phenol-like Breath Marker: The Future of Induced Volatolomics in Cancer Risk Pre-warning"

### Test Report

| | |
|---|---|
| Sample Name | Phenyl-β-D-glucuronide (CAS: 17685-05-1) |
| Client | Shanghai Jiao Tong Universty |
| Report Number | FT-20230204002-En-1 |

**Shanghai Fuda Testing Technology Group CO., Ltd.**

Second Floor, Fuhua Building Fudan University, No. 525, Guoquan Road, Yangpu District, Shanghai, China.
Service Hotline：021-61996230

| | | | |
|---|---|---|---|
| Sample Name | Phenyl-β-D-glucuronide(CAS: 17685-05-1) | | |
| Sample Quantity | 1 | Sample Batch | / |
| Sample Status | Intact | Sample Number | FT230204002 |
| Client | Shanghai Jiao Tong Universty | | |
| Communication Information of Client | No. 800 Fenghua Road, Minghang District, Shanghai, P.R. China | | |
| Test Category | Commission Test | | |
| Sample Arrival Date | 2023.03.29 | | |
| Test Cycle | 2023.03.29-2023.04.15 | | |
| Standards and Methods | Please refer to next page(s). | | |
| Test Results | This report only provides the measured values. See the summary page of test results in this report for details. | | |
| Remarks | —— | | |

Drafter：______ Signer：______

Reviewer：______ Issued Date：2024-01-29

### Test Result(s):

The extract of the test article was evaluated for its potential to induce skin sensitization in the Guinea Pig Maximization Test.

The test article were extracted with 0.9% sodium chloride injection. The test article extract was ed into guinea pigs and applied topically for induction. Five control animals were treated accordingly but with the negative control alone. During the recovery period, the extract of the test article was used for challenge test.

Under the conditions of this test, the extract of the test article did not cause skin sensitization reaction, and the sensitization positive rate was 0%

The study was carried out in accordance with the standard operating procedure. The test process was conducted in compliance with the requirements of CNAS-CL01:2018 (ISO/IEC17025:2017, IDT) and RB/T214-2017.

**1.0 Purpose**

The test was designed to evaluate the potential of a test article to cause skin sensitization using Guinea Pig

Maximization Test.

**3.0 Test and control articles**

3.1 Test article (The information about the test article was supplied by the sponsor wherever applicable.)

Test article name: Phenyl-β-D-glucuronide (CAS: 17685-05-1)

Sterilization state: Non-sterile

Model: 0.4 mg/ml

Size: 20ml

Lot/ Batch#: 20230325

Physical State: Liquid

Color: See the photo

Density: N/S

Stability: Stable

Solubility: Dissoluble

Test Article Material: Solution

Packing Material: Glass bottle

Storage Condition: Room Temperature

3.2 Control Article

Name: 0.9% sodium chloride injection (SC)

Manufacturer: Anhui Fengyuan Pharmaceutical Co., Ltd. Huaihai Pharmaceutical Factory

Size: 500 ml

Physical State: Liquid

Color: Colourless

Lot/ Batch#: 220923102

Storage Condition: Room Temperature

**4.0 Identification of test system**

Species: White Guinea Pig

Number: 15 (10 for test group and 5 for control group)

Sex: Males

Health status: Healthy, not previously used in other experimental procedures

Housing: Animals were housed in groups in cages identified by a card indicating the lab number and test code.

Animal identification: Animal marker pen

The quarantine period: 3 days

**5.0 Animal Care and Maintenance**

Animal purchase: Provided by Danyang Changyi experimental animal breeding Co., Ltd <Permit Code: SCXK (SU) 2021-0002>

Bedding: NA

Feed: Guinea Pig Diet, Beijing Keao Xieli Feed Co., Ltd.

Water: Drinking water met the Standards for Drinking Water Quality GB 5749-2006

Cages: Plastic cage, Suzhou Fengqiao purification equipment Co.,Ltd.

Environment: Temperature 18-29°C, Relative humidity 40%-70%, Lights 12 hours light/dark cycle

Personnel: Associates involved were appropriately qualified and trained

Selection: Only healthy, previously unused animals were selected

Veterinarian: Vet takes care of the whole course

Ethics: Test methods of operation were reviewed and approved by the Commission on Science Standard animal ethics

There were no known contaminants present in the feed, water, or bedding expected to interfere with the test data.

**6.0 Justification of the test system**

6.1 The guinea pig is believed to be the most sensitive animal model for this type of study. The susceptibility of the guinea pig to a known sensitizing agent, dinitrochlorobenzene (DNCB) has been substantiated at SSMT.

6.2 The test article was extracted and administered in vivo through a medium compatible with the test system, which is considered as the best route of administration.

**7.0 Instruments and reagents**

7.1 Instruments

Electronic balance (SSMT-494)

Electronic balance (SSMT-532)

Electronic balance (SSMT-147)

Clean bench (SSMT-501)

Thermostatic oscillation incubator(SSMT-564)

Temperature recorder(SSMT-849)

7.2 Reagents

Sodium dodecyl sulfate (SDS)

Freund's Adjuvant, Complete liquid

**8.0 Experiment design and dose**

8.1 Sample preparation

The test article was extracted as Table 1. Extract was checked and used immediately after extraction without the process of filtering, centrifugation, dilution, etc. The pH of the extract was not adjusted prior to testing. The preparation process was aseptic. The control article was prepared under the same condition.

Table 1 Sample Preparation

| Aseptic Sampling | | | Aseptic Agitation Extraction In Inert Container | | | | Final Extract |
|---|---|---|---|---|---|---|---|
| Sampling Manner | Test phase | Actually Sampling | Extraction solvent | Extraction ratio | Solvent volume | Condition | Clear or Not |
| Random sampling | Intradermal induction phase | 1.39 g | 0.9% sodium chloride injection | 0. 1 g : 1 ml | 13.9 ml | 37℃ , 72 h | Clear |
| | Topical induction phase | 1.70 g | | | 17.0 ml | 37℃ , 72 h | Clear |
| | Challenge phase | 1.73 g | | | 17.3 ml | 37℃ , 72 h | Clear |

8.2 Test method

8.2.1 Intradermal induction phase

A pair of 0.1 ml intradermal injections was made for each animal, at the sites (A, B and C) in the clipped intrascapular region as shown in the following Figure 1

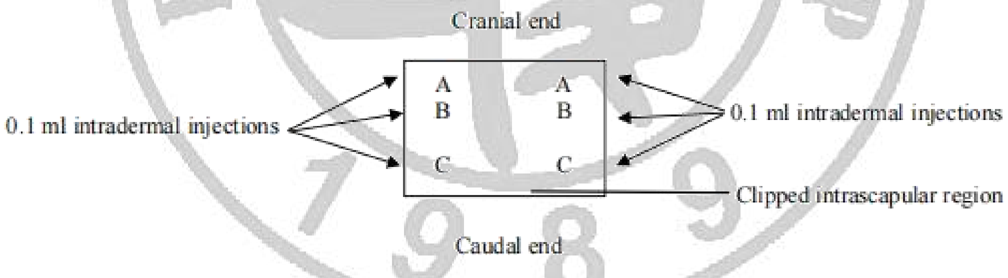


Figure 1 Location of intradermal injection sites

Site A: A 50:50 volume ratio stable emulsion of Freund’s complete adjuvant mixed with the solvent.

Site B: The test sample (undiluted extract); inject the control animals with the control articles alone.

Site C: The test sample at the concentration used at site B, emulsified in a 50:50 volume ratio stable emulsion of Freund's complete adjuvant and the solvent (50%); inject the control animals with an emulsion of the blank liquid with adjuvant.

8.2.2 Topical induction phase

At 7 d after the intradermal induction phase, administer the test sample by topical application to the intrascapular region of each animal, using a patch of area approximately 8 cm2 (absorbent gauze) so as to cover the intradermal injection sites. Use the concentration selected in site B of intradermal induction phase. The concentration in Intradermal induction phase did not produce irritation, pretreated the area with 10% sodium dodecyl sulfate massaged into the skin 24 hours before the patch is applied. Secure the patches with an occlusive dressing. Remove the dressings and patches after 48 h.

Treat the control animals similarly, using the blank liquid alone.

8.2.3 Challenge phase

At 15 d after completion of the topical induction phase, challenge all test and control animals with the test sample. Administer the test sample and a blank by topical application to left and right abdomen of animals respectively, using absorbent gauze (about 8 cm2) soaked with 0.5ml extracts or solvent control. Secure with an occlusive dressing. Remove the dressings and patches after 24 h.

8.3 Observation of animal

Observe the appearance of the challenge skin sites of the test and control animals 24 h and 48 h after removal of the dressings. Full-spectrum lighting was used to visualize the skin reactions. Describe and grade the skin reactions for erythema and oedema according to the Magnusson and Kligman grading given in Table 2 for each challenge site and at each time interval.

Table 2 Magnusson and Kligman scale

| Patch test reaction | Grading scale |
|---|---|
| No visible change | 0 |
| Discrete or patchy erythema | 1 |
| Moderate and confluent erythema | 2 |
| Intense erythema and/or swelling | 3 |

**9.0 Evaluation criteria**

Magnusson and Kligman grades of 1 or greater in the test group generally indicate sensitization, provided grades of less than 1 are seen in control animals.

If grades of 1 or greater are noted in control animals, then the reactions of test animals which exceed the most severe reaction in control animals are presumed to be due to sensitization.

If the response is equivocal, rechallenge is recommended to confirm the results from the first challenge.

**10.0 Results of the test**

The skin response of guinea pigs is shown in Table 3. The positive control test results are in shown Table 4.

Table 3 Guinea pig Sensitization Dermal Reactions

| Group | Animal Number | Excitation patch removed 24 h | Excitation patch removed 48h | Positive rate after challenge phase | Weight range before injection (g) | Weight range after experiment (g) | Abnormal appearance except dermal reactions |
|---|---|---|---|---|---|---|---|
| Control | 2023-01174-01-J1001 | 0 | 0 | 0% | 321.0-346.7 | 461.9-495.4 | None |

| Group | Animal Number | Excitation patch removed 24 h | Excitation patch removed 48h | Positive rate after challenge phase | Weight range before injection (g) | Weight range after experiment (g) | Abnormal appearance except dermal reactions |
|---|---|---|---|---|---|---|---|
| Control | 2023-01174-01-J1002 | 0 | 0 | 0% | 321.0-346.7 | 461.9-495.4 | None |
| | 2023-01174-01-J1003 | 0 | 0 | | | | None |
| | 2023-01174-01-J1004 | 0 | 0 | | | | None |
| | 2023-01174-01-J1005 | 0 | 0 | | | | None |
| Test | 2023-01174-01-J2001 | 0 | 0 | 0% | 330.6-368.4 | 482.5-510.5 | None |
| | 2023-01174-01-J2002 | 0 | 0 | | | | None |
| | 2023-01174-01-J2003 | 0 | 0 | | | | None |
| | 2023-01174-01-J2004 | 0 | 0 | | | | None |
| | 2023-01174-01-J2005 | 0 | 0 | | | | None |
| | 2023-01174-01-J2006 | 0 | 0 | | | | None |
| | 2023-01174-01-J2007 | 0 | 0 | | | | None |
| | 2023-01174-01-J2008 | 0 | 0 | | | | None |
| | 2023-01174-01-J2009 | 0 | 0 | | | | None |
| | 2023-01174-01-J2010 | 0 | 0 | | | | None |

Table 4 Results of positive control test

| Group | Animal Number | Excitation patch removed 24 h | Excitation patch removed 48 h | Positive rate after challenge phase | Weight range before injection (g) | Weight range after experiment (g) | Abnormal appearance except dermal reactions |
|---|---|---|---|---|---|---|---|
| | 2023-01521-01-X1001 | 0 | 0 | | | | None |
| | 2023-01521-01-X1002 | 0 | 0 | | | | None |
| Control | 2023-01521-01-X1003 | 0 | 0 | 0% | 319.8-362.4 | 468.7-524.5 | None |
| | 2023-01521-01-X1004 | 0 | 0 | | | | None |
| | 2023-01521-01-X1005 | 0 | 0 | | | | None |
| | 2023-01521-01-X2001 | 3 | 2 | | | | None |
| | 2023-01521-01-X2002 | 2 | 2 | | | | None |
| Test | 2023-01521-01-X2003 | 2 | 1 | 100% | 326.8-358.6 | 460.3-491.2 | None |
| | 2023-01521-01-X2004 | 3 | 2 | | | | None |
| | 2023-01521-01-X2005 | 3 | 3 | | | | None |

Note: The skin sensitized positive control test is conducted every three months. The data was from the report SSMT-R-2023-01521-01 ( Date: 2023.03.13-2023.04.09) and the allergenic rate is 100%(Positive sample: DNCB, Intradermal induction phase : 0. 1%; Topical induction phase: 0.5%; Challen e phase: 0. 1%).

Under the conditions of this study, the test article did not show significant evidence of causing skin sensitization in the guinea pig. The skin sensitization rate was determined with 0%.

***The End of the Report***

Sample Photo:

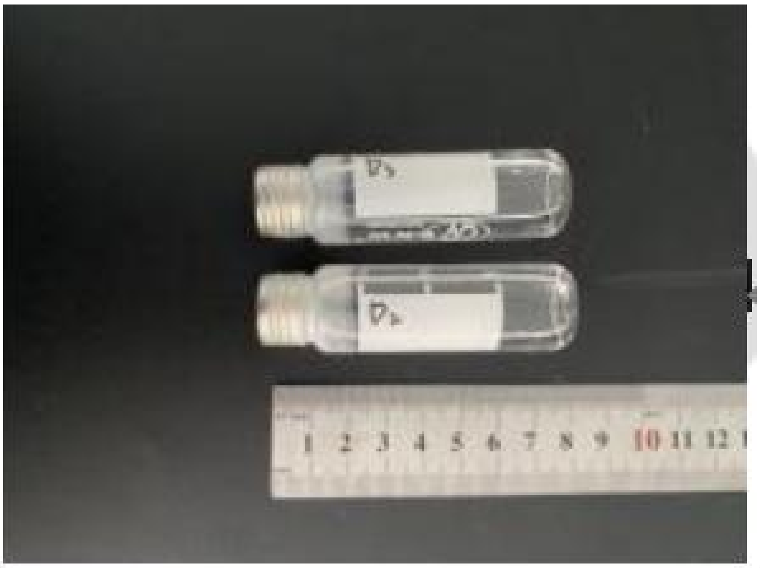


### Additional Instructions of the Report

1. The Report would be invalid without "Special Seal for Report of Shanghai Fuda Testing Technology Group CO., Ltd.".
2. Any institution is not permitted to duplicate the report, if needed please submit a formal application.
3. Any objection to the report should be interposed in 10 days from the date of report is issued. Overdue would not be admissible.
4. The report is only responsible for the sample provided by the applicant. The sample will be kept for30 days after the date of report is issued.
5. The company shall perform the duty of confidentiality to the technical documents, report, contract documents and other business secrets of the applicant.
6. When the report is not stamped with the qualification identification mark (CMA), it indicates that the relevant projects have not obtained the qualification identification. The data and results are only used for scientific research, teaching and internal quality control, not for social justice. The Chinese version shall prevail.
