## Supplementary reports for the maximization dosage test for "Discovery of Phenyl-β-D-glucuronide Medical Function for in Vivo Producing Handheld Gas Sensor Detectable Phenol-like Breath Marker: The Future of Induced Volatolomics in Cancer Risk Pre-warning"

### Test Report

| | |
|---|---|
| Sample Name | Phenyl-β-D-glucuronide (CAS: 17685-05-1) |
| Client | Shanghai Jiao Tong Universty |
| Report Number | FT-20230204002-En-2 |

**Shanghai Fuda Testing Technology Group CO., Ltd.**

Second Floor, Fuhua Building Fudan University, No. 525, Guoquan Road, Yangpu District, Shanghai, China.
Service Hotline：021-61996230

| | | | |
|---|---|---|---|
| Sample Name | Phenyl-β-D-glucuronide(CAS: 17685-05-1) | | |
| Sample Quantity | 1 | Sample Batch | / |
| Sample Status | Intact | Sample Number | FT230204002 |
| Client | Shanghai Jiao Tong Universty | | |
| Communication Information of Client | No. 800 Fenghua Road, Minghang District, Shanghai, P.R. China | | |
| Test Category | Commission Test | | |
| Sample Arrival Date | 2023.03.29 | | |
| Test Cycle | 2023.03.29-2023.04.15 | | |
| Standards and Methods | Please refer to next page(s). | | |
| Test Results | This report only provides the measured values. See the summary page of test results in this report for details. | | |
| Remarks | —— | | |

Drafter：__________ Signer：__________

Reviewer：__________ Issued Date：2024-01-29

### Test Result(s):

The test article was evaluated for acute systemic toxicity after the extract of the test article being injected into white mice.

The test articles were respectively extracted with 0.9% sodium chloride injection. The extract and negative control were injected into the tail vein in polar test group.

Within the monitoring period, the testing group exhibited the same response as control group.

The results indicated that the extract of the test article did not lead the mice to toxicosis under the test condition of this study.

The study was carried out in accordance with the standard operating procedure. The test process was conducted in compliance with the requirements of CNAS-CL01:2018 (ISO/IEC17025:2017, IDT) and RB/T214-2017

#### 1.0 Purpose

The test was designed to evaluate the potential of a test article to cause acute systemic toxicity.

#### 3.0 Test and control articles

3.1 Test article (The information about the test article was supplied by the sponsor wherever applicable.) Test article name: Phenyl-β-D-glucuronide (CAS: 17685-05-1)

Sterilization state: Non-sterile

Model: 0.4 mg/ml

Size: 20ml

Lot/ Batch#: 20230325

Physical State: Liquid

Color: See the photo

Density: N/S

Stability: Stable

Solubility: Dissoluble

Test Article Material: Solution

Packing Material: Glass bottle

Storage Condition: Room Temperature

3.2 Control Articles

3.2.1 Polar negative control

Name: 0.9% Sodium chloride injection (SC)

Manufacturer: Anhui Fengyuan Pharmaceutical Co., Ltd. Huaihai Pharmaceutical Factory

Size: 500 ml

Physical State: Liquid

Color: Colourless

Lot/ Batch#: 220923102

Storage Condition: Room Temperature

**4.0 Identification of test system**

Species: ICR mouse

Number: 10 for polar test group(5 for test and 5 for control)

Sex: males

Weight: 17-23 g. The weight variation of animals used within a sex did not exceed ±20 % of the mean weight.

Health status: Healthy young adult

Housing: Reared in groups and each cage was identified by a card indicating the lab number and test code.

Animal identification: Animal marker pen

The quarantine period: 5 days

#### 5.0 Animal Care and Maintenance

Animal purchase: Provided by Hangzhou Qizhen Experimental Animal Technology Co., Ltd. (Production) <Permit Code: SCXK (ZHE) 2022-0005>

Environment: Temperature 20-26°C, Relative humidity 40%-70%, Lights 12 hours light/dark cycle

Personnel: Associates involved were appropriately qualified and trained

Selection: Only healthy, previously unused animals were selected

Veterinarian: Vet takes care of the whole course

Ethics: Test methods of operation were reviewed and approved by the Commission on Science Standard animal ethics.

There were no known contaminants present in the feed, water expected to interfere with the test data.

#### 6.0 Justification of the test system

6.1 The mice were used in this study because they have historically been used in safety evaluation studies and the guidelines have no alternative (non-animal) method. The species and number of animals as well as the route of administration used are recommended by Standard guidelines.

6.2 The test article was exposed to the test system through a solvent compatible with the test system. This was the

optimal route of administration available in this test system.

#### 7.0 Instruments

Thermostatic oscillation incubator (SSMT-564)

Temperature recorder (SSMT-849)

Clean bench (SSMT-501)

Electronic balance (SSMT-493)

Electronic balance (SSMT-147)

#### 8.0 Experiment design and dose

8.1 Sample preparationThe test article was extracted as Table 1. Extract was checked and used immediately after extraction without the process of filtering, centrifugation, dilution, etc. The pH of the extract was not adjusted prior to testing. The preparation process was aseptic. The control article was prepared under the same condition.

Table 1 Sample Preparation

| Aseptic Sampling | | Aseptic Agitation Extraction In Inert Container | | | | Final Extract |
|---|---|---|---|---|---|---|
| Sampling Manner | Actually Sampling | Extraction solvent | Extraction ratio | Solvent volume | Condition | Clear or Not |
| Random sampling | 2.05 g | 0.9% sodium chloride injection | 0.1 g : 1 ml | 20.5 ml | 37 °C, 72 h | Clear |

8.2 Test method

Take 10 mice for polar test group.(5 for test group and 5 for control group).

Polar test group: Fix the mouse and inject the test article extract or negative control into its tail vein.

The injection dosage is 50 ml/kg.

8.3 Observation of animal

After injection, observe the instant response of the mouse. Further more, monitor and record the status of testing group and control group (e.g. general state, toxicosis symptom, mortality) in 4 h, 24 h, 48 h and 72 h. And animals were weighed and recorded at 24 h, 48 h and 72 h.

**9.0 Evaluation criteria**

9.1 If during the observation period of an acute systemic toxicity test none of the animals treated with the test article shows a significantly greater biological reactivity than animals treated with the vehicle control, the sample meets the requirements of this test.

9.2 Using five animals, if two or more animals die, or if behavior such as convulsions or prostration occurs in two or more animals, or if a body weight loss greater than 10 % occurs in three or more animals, the sample does not meet the requirements of the test.

9.3 If any animals treated with the sample show only slight signs of biological reactivity, and not more than one animal shows gross symptoms of biological reactivity or dies, repeat the testing using groups of ten animals.

9.4 On the repeat test, if all ten animals treated with the sample show no scientifically meaningful biological reactivity above the vehicle control animals during the observation period, the sample meets the requirements of this test.

**10.0 Results of the test**

Within the monitoring period, the testing group exhibited the same response as control group. See Table 2 and Table 3.

Table 2 Animal Weight and Dose

| Extraction solvent | Group | Animal No. | 0 h Weight (g) | Dose (ml) | Weight (g) | | |
|---|---|---|---|---|---|---|---|
| | | | | | 24 h | 48 h | 72 h |
| 0.9% sodium chloride injection | Negative Control | 2023-01174-02-J1001 | 18.2 | 1.0 | 19.1 | 20.5 | 21.4 |
| | | 2023-01174-02-J1002 | 21.6 | 1.1 | 22.6 | 24.0 | 25.4 |
| | | 2023-01174-02-J1003 | 19.1 | 1.0 | 20.3 | 21.4 | 22.9 |
| | | 2023-01174-02-J1004 | 18.3 | 1.0 | 19.7 | 21.0 | 22.4 |
| | | 2023-01174-02-J1005 | 19.1 | 1.0 | 20.4 | 21.3 | 22.5 |
| | Test Article Group | 2023-01174-02-J2001 | 19.0 | 1.0 | 20.2 | 21.6 | 22.7 |
| | | 2023-01174-02-J2002 | 21.0 | 1.1 | 22.2 | 23.5 | 24.8 |
| | | 2023-01174-02-J2003 | 21.2 | 1.1 | 22.6 | 23.9 | 25.3 |
| | | 2023-01174-02-J2004 | 19.7 | 1.0 | 21.0 | 22.4 | 23.8 |
| | | 2023-01174-02-J2005 | 22.0 | 1.1 | 23.0 | 23.8 | 24.6 |

Table 3 Observed Signs

| Extraction solvent | Group | Animal No. | Observation | | | |
|---|---|---|---|---|---|---|
| | | | 4 h | 24 h | 48 h | 72 h |
| 0.9% sodium chloride injection | Negative Control | 2023-01174-02-J1001 | - | - | - | - |
| | | 2023-01174-02-J1002 | - | - | - | - |
| | | 2023-01174-02-J1003 | - | - | - | - |
| | | 2023-01174-02-J1004 | - | - | - | - |
| | | 2023-01174-02-J1005 | - | - | - | - |
| | Test Article Group | 2023-01174-02-J2001 | - | - | - | - |
| | | 2023-01174-02-J2002 | - | - | - | - |
| | | 2023-01174-02-J2003 | - | - | - | - |
| | | 2023-01174-02-J2004 | - | - | - | - |
| | | 2023-01174-02-J2005 | - | - | - | - |

Note:-shows there were no signs of toxicity

Under the conditions of this study, the extract of test article showed no significant evidence of causing acute systemic toxicity in the mice in polar test group

***The End of the Report***

#### Sample Photo:

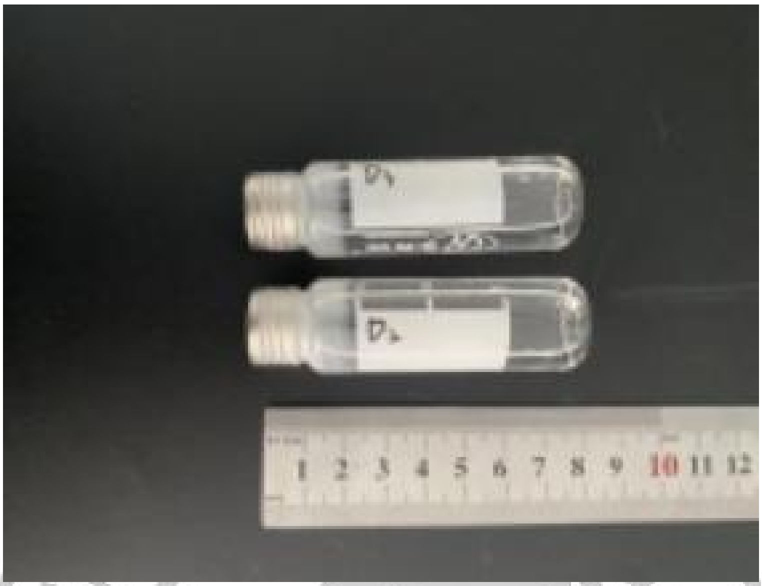


### Additional Instructions of the Report

1. The Report would be invalid without "Special Seal for Report of Shanghai Fuda Testing Technology Group CO., Ltd.".
2. Any institution is not permitted to duplicate the report, if needed please submit a formal application.
3. Any objection to the report should be interposed in 10 days from the date of report is issued. Overdue would not be admissible.
4. The report is only responsible for the sample provided by the applicant. The sample will be kept for30 days after the date of report is issued.
5. The company shall perform the duty of confidentiality to the technical documents, report, contract documents and other business secrets of the applicant.
6. When the report is not stamped with the qualification identification mark (CMA), it indicates that the relevant projects have not obtained the qualification identification. The data and results are only used for scientific research, teaching and internal quality control, not for social justice. The Chinese version shall prevail.
